# Tuft cells transdifferentiate to neural-like progenitor cells in the progression of pancreatic cancer

**DOI:** 10.1101/2024.02.12.579982

**Authors:** Daniel J. Salas-Escabillas, Megan T. Hoffman, Jacee S. Moore, Sydney M. Brender, Hui-Ju Wen, Simone Benitz, Erick T. Davis, Dan Long, Allison M. Wombwell, Nina G. Steele, Rosalie C. Sears, Ichiro Matsumoto, Kathleen E. DelGiorno, Howard C. Crawford

## Abstract

Pancreatic ductal adenocarcinoma (PDA) is partly initiated through the transdifferentiation of acinar cells to metaplastic ducts that act as precursors of neoplasia and cancer. Tuft cells are solitary chemosensory cells not found in the normal pancreas but arise in metaplasia and neoplasia, diminishing as neoplastic lesions progress to carcinoma. Metaplastic tuft cells (mTCs) function to suppress tumor progression through communication with the tumor microenvironment, but their fate during progression is unknown. To determine the fate of mTCs during PDA progression, we have created a lineage tracing model that uses a tamoxifen-inducible tuft-cell specific Pou2f3^CreERT/+^ driver to induce transgene expression, including the lineage tracer tdTomato or the oncogene *Myc*. mTC lineage trace models of pancreatic neoplasia and carcinoma were used to follow mTC fate. We found that mTCs, in the carcinoma model, transdifferentiate into neural-like progenitor cells (NRPs), a cell type associated with poor survival in PDA patients. Using conditional knock-out and overexpression systems, we found that *Myc* activity in mTCs is necessary and sufficient to induce this Tuft-to-Neuroendocrine-Transition (TNT).

## Introduction

Pancreatic ductal adenocarcinoma (PDA), the most common form of pancreatic cancer, is a devastating disease with a 5-year survival rate of only 13% ^1^. Effective treatment options are few, and reliable early detection methods needed to enhance treatment effectiveness are non-existent. PDA is initiated by oncogenic mutations in the *KRAS* gene, causing irreversible transdifferentiation of normal acinar cells into metaplastic ducts. Metaplastic ducts then progress to pancreatic intraepithelial neoplasia (PanIN) ^2^ and, with the acquisition of mutations in tumor suppressor genes such as *TP53*, *CDKN2A*, or *SMAD4 ^3, 4^*, progress to invasive carcinoma. Metaplastic ducts are a heterogeneous population of reactive epithelial cells ^5^ comprised of at least 6 distinct identities: gastric neck cells ^6^, gastric pit-like cells, gastric chief-like cells, ductal senescent cells, metaplastic neuroendocrine cells (mNECs), and specialized chemosensory cells known as metaplastic tuft cells (mTCs) ^7^.

Normal tuft cells are solitary chemosensory cells found throughout the body that sense and respond to environmental cues to maintain tissue homeostasis. Functionally, they use the canonical gustatory signal transduction pathway, with specific stimulants and responses depending on context and the organ in which they reside ^8–10^. In pancreatic neoplasia, metaplastic tuft cells (hereafter referred to as mTCs) have been proposed to be a class of tumor progenitor cells ^9, 11^. Still, functionally, they suppress progression to PDA via inhibition of anti-inflammatory immune cells through crosstalk with the tumor microenvironment ^8, 10^. mTCs are identified by staining for markers such as Acetylated-*Tuba1a^10^*, *Dclk1*, *Vav1*, *Cox1^9^*, *Cox2^12–14^*, *Trpm5 ^15^*, and *Pou2f3*, the latter being the master transcriptional regulator of tuft cell genesis ^15, 16^. mTCs are not typically found in the normal pancreas and only appear after prolonged damage, such as in chronic pancreatitis ^17^ or the initiation of neoplasia ^9, 11^, arising from the transdifferentiation of acinar cells. As neoplasia progresses to invasive carcinoma, the mTC population gradually disappears ^9, 11^, challenging their putative role as classic tumor progenitor cells.

In the current study, to address the hypothesis that mTCs act as tumor progenitor cells in pancreatic cancer, we lineage-traced their fate during disease progression in mouse models of pancreatic disease. To accomplish this, we used a dual recombinase model wherein tumorigenesis is initiated by the expression of FlpO recombinase knocked into the pancreas-specific *Ptf1a* locus, driving the expression of an Frt-STOP-Frt oncogenic *Kras^G12D/+^*allele and the deletion an Frt-flanked allele of *Trp53 ^18^*. This model allowed us to use an inducible *Pou2f3^CreERT/+^* driver ^19^ to target the expression of a *ROSA26^LSL-TdTomato^*lineage trace reporter specifically to mTCs ^16, 20^ in both neoplasia and carcinoma, indelibly labeling mTCs and their progeny. In the *Ptf1a^FlpO/+^; Kras^G12D^*neoplasia model, mTCs and other metaplastic cellular subtypes, including neuroendocrine-like enteroendocrine cells ^7, 21, 22^ arise from acinar cells, independent from one another. However, in the *Ptf1a^FlpO/+^; Kras^FSF- G12D/+^; Trp53^Frt-Exons 2 to 5-Frt/+^* carcinoma model, we found that acinar cell-derived mTCs transdifferentiate to neural-like progenitor cells (NRPs) ^21^, also known as ductal neuroendocrine cells ^23^. NRPs in PDA patients correlate with worse survival, are resistant to chemotherapy, and their genesis is associated with elevated *MYC* activity ^21, 23^. Indeed, modulating the expression of *Myc* to mTCs demonstrated that *Myc* activity was necessary and sufficient to drive the transdifferentiation of mTCs to NRPs.

## RESULTS

### Pou2f3-CreERT specifically targets recombination in metaplastic tuft cells in the pancreas

Tuft cells are not found in the normal mouse pancreas but arise from the transdifferentiation of acinar cells in conditions of tissue damage and early neoplasia ^24^. As neoplasia progresses, metaplastic tuft cells (mTCs) decrease in number and are absent in carcinoma ^9, 11, 24^. To determine the fate of mTCs during neoplastic progression, we developed a novel dual recombinase lineage trace system, wherein Ptf1a^Flpo/+ 25^ initiates expression of oncogenic Kras from a Kras^Frt-Stop-Frt-G12D^ allele (*KF*) to drive neoplasia ^18^ (Fig. 1A, 1D). When combined with a conditional knock-out trp53 allele with exons 2-5 flanked by Frt recombination sites (Trp53^Frt-Exons 2 to 5-Frt/+^), lesions rapidly progress from neoplasia to carcinoma (*KPF*) (Fig. 1B, 1D). Within this system, we introduced a Cre^ERT^ driver knocked into the *Pou2f3* locus ^19^, which encodes the transcription factor master regulator of tuft cell genesis ^15, 16, 20, 26, 27^. *Pou2f3^CreERT/+^* was used to drive recombination and expression of a *ROSA26^LSL-tdTomato^* reporter (tdTom) ^28^ in a tamoxifen-inducible manner (P2f3T) ^8, 20, 27^, allowing us to lineage trace mTCs temporally in our neoplasia (*KF-P2f3T)* and carcinoma (*KPF-P2f3T)* models.

**Figure 1.**
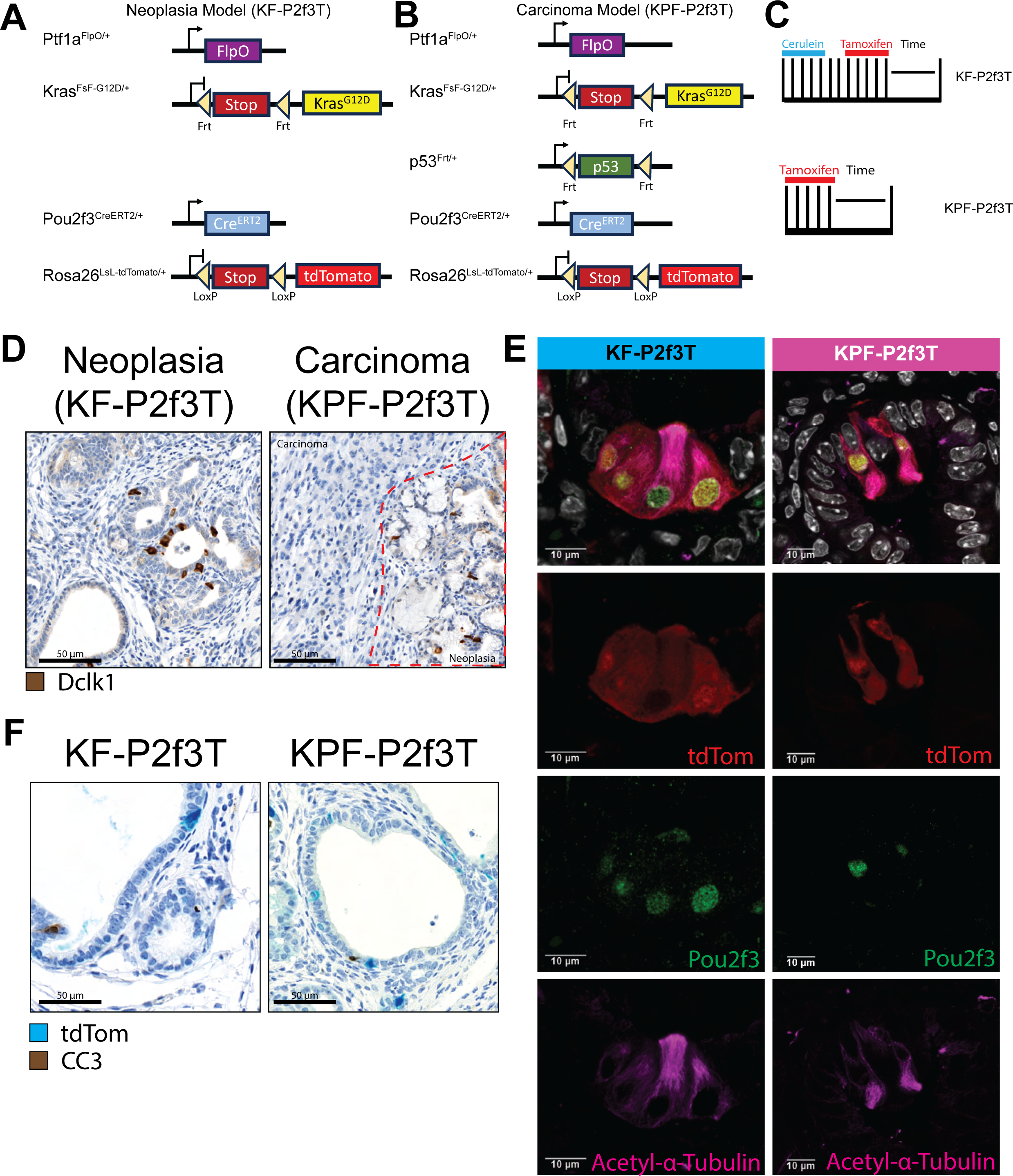
Lineage trace model shows tuft cell specific expression of tdTomato. Genetic strategy to trace the fate of metaplastic tuft cells (mTCs) in pancreatic cancer progression using a dual recombinase neoplasia (*KF-P2f3T*) model **(A)** and carcinoma (*KPF-P2f3T*) model **(B)**. **C)** Treatment regimen for neoplasia and carcinoma Models. The models have both pancreatic progression and reporter specifically in tuft cells. The *KF-P2f3T* model utilizes Cerulein once daily, followed by 2 weeks recovery, then once daily tamoxifen treatment for 5 days. The *KPF-P2f3T* model is aggressive as it contains one null p53 allele. Cerulein is not necessary and so only Tamoxifen is treated once daily for 5 days. Mice are allowed to age for different time points to determine lineage of metaplastic tuft cells in progression of pancreatic cancer. **D)** Dclk1 in Brown identifies tuft cells in neoplasia model (left) and carcinoma model (right). **E)** Specific expression of tdTomato reporter in both models. White represents Hoechst staining to mark the nucleus, tdTomato in red to show recombination of tdTomato (tdTom), Pou2f3, the master regulator of tuft cells in Green, and Acetylated alpha tubulin in Magenta to mark filamentous actin in the tufts of the mTCs. **F)** Dual IHC of tdTom (Teal) and CC3 (Brown) stained *KF-P2f3T* and *KPF-P2f3T* mice to determine cell death of mTCs in neoplasia and carcinoma. Black arrows indicate tdTom+ cells in lesions. Scale bars = 10 μm unless otherwise noted.

To determine the fidelity of our systems in labeling tuft cells, we treated *KF-P2f3T* with cerulein at 8-9 weeks of age to induce mild pancreatitis and accelerate transformation, then allowed the mice to recover for 3 weeks to optimize mTC genesis ^9, 11^. The *KPF-P2f3T* carcinoma model induced sufficient transformation and mTC formation at 11-12 weeks of age with no additional treatment (Fig. 1C). Cerulein-treated *KF-P2f3T* and 11–12-week-old *KPF-P2f3T* mice were treated with tamoxifen to induce tdTom reporter activity, and the pancreata harvested 1 day later. We co-stained tissues for tdTom and the tuft cell markers Pou2f3, Acetylated-α-tubulin, Cox1, Dclk1, Trpm5, and Vav1 (Fig. 1E and Supp. Fig. 1A, 1B) by immunofluorescence (IF). In both models, we found co-expression of tdTom exclusively in cells expressing mTC markers (Supp. Fig. 1A, 1B), confirming the system’s specificity and allowing us to trace mTC fate throughout the progression of PDA.

### Metaplastic tuft cells change identity in pancreatic adenocarcinoma

To determine the fate of the mTC population over time, we collected tissue from our *KF-P2f3T* and *KPF-P2f3T* mice 3 days and 7 weeks post-tamoxifen. In the *KF-P2f3T* neoplasia model, all tdTom^+^ cells co-expressed tuft cell markers (Fig. 2A). The population of mTCs dropped off slightly over this period but largely remained stable (Fig. 2B, *Black*). In contrast, in the *KPF-P2f3T* carcinoma model, the mTC population dropped drastically by 7 weeks, consistent with their disappearance as PanIN progresses to PDA ^9, 11^ (Fig. 2A and 2B, *Black*). We found no evidence that mTC attrition was associated with apoptosis by co-staining for cleaved caspase 3 (CC3) (Fig. 1F). Instead, we noted that tdTom^+^ cells persisted in this model but no longer expressed mTC markers (Fig. 2A), suggesting that mTCs take on a new cellular identity.

**Figure 2.**
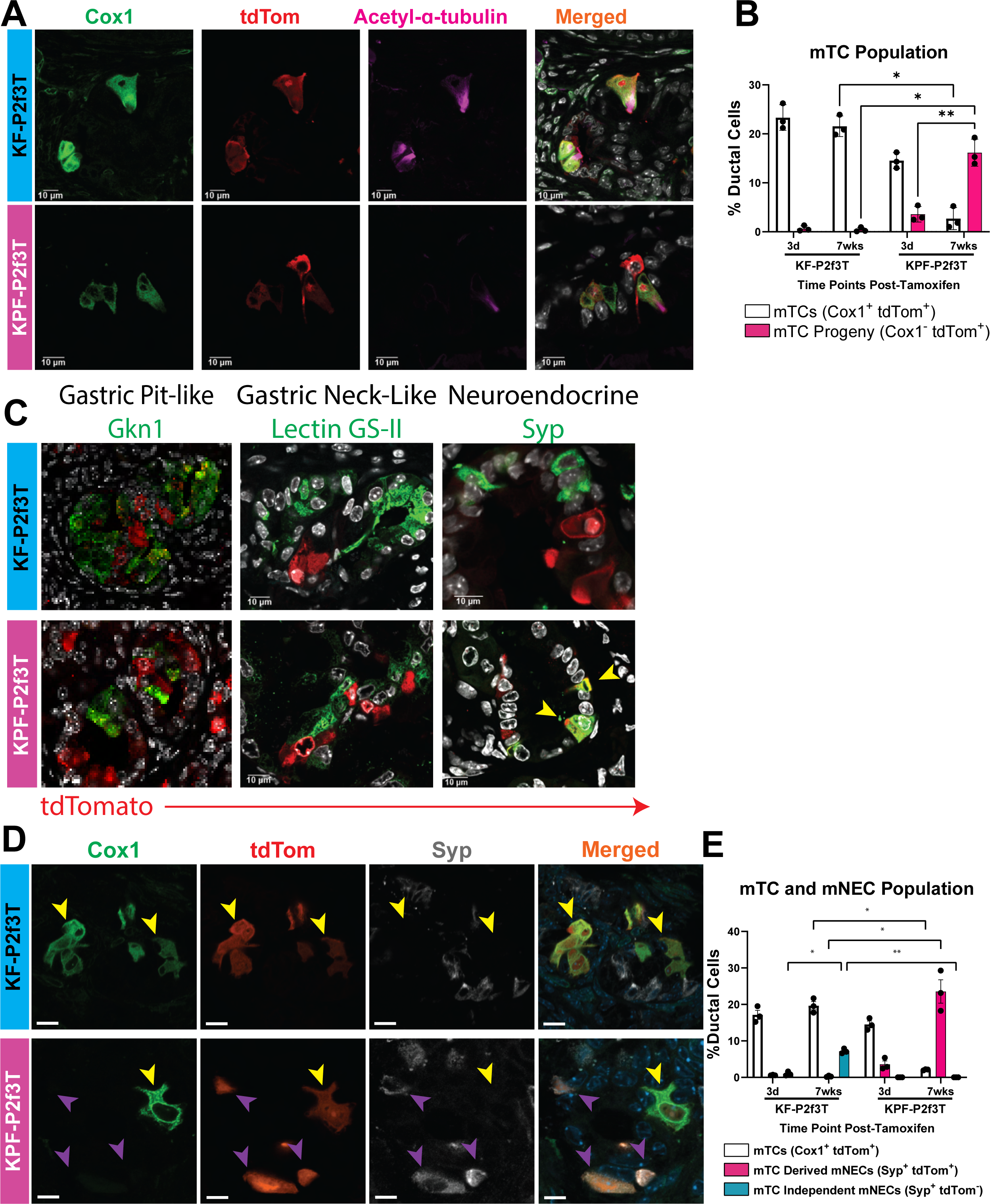
Lineage tracing mTCs reveals transdifferentiation to metaplastic neuroendocrine cells (mNECs). Dual recombinase models show fate of mTCs in progression of pancreatic cancer. **A)** Mice harvested 3 days post-tamoxifen treatment for labeling mTCs. Hoechst stains nucleus of cells (White), Cox1 is used to identify mTCs (Green), tdTom reporter to determine fate of mTCs (Red) and acetylated-α-tubulin (Magenta) marks filamentous actin on tuft of mTCs. **B)** Quantification of Percentage of mTC (White) and mTC Progeny (Pink) among ductal cells at different time points post-tamoxifen treatment in *KF-P2f3T* and *KPF-P2f3T* models. mTCs are identified by Cox1 and tdTom, mTC Progeny are only tdTom positive. (n = 3/group) p-values were calculated using a two-way ANOVA on Prism GraphPad 7. * = p < 0.05, ** = p < 0.01. **C)** Co-staining of tdTomato (Red) expression with remaining metaplastic ductal cell subsets. In Green, Gkn1, Lectin GS-II, and Synaptophysin (Syp) mark gastric pit-like, gastric neck-like, and neuroendocrine cells respectively in pancreatic metaplastic ducts. **D)** 7 weeks post-tamoxifen labeling of neoplasia and carcinoma models exposes shift of populations. Cox1 (Green) marks mTCs, tdTomato (Red) identifies reporter, Syp (White) marks neuroendocrine cells, and Blue represents DAPI Staining to identify the nucleus of a cell. **E)** Quantification of percentage of mTCs (Cox1+ tdTom+) in White, mTC-derived mNECs (Syp+ tdTom+) in Pink, mTC-independent mNECs (Syp+ tdTom-) in Green. (n = 3/group). p-values were calculated using a two-way ANOVA on Prism GraphPad 7. * = p < 0.05, ** = p < 0.01. Scale bars = 10 μm unless otherwise noted.

Quantitation of the tdTom expression throughout multiple pancreata revealed that, in the *KPF-P2f3T* carcinoma model, the Cox1^+^ mTC population dropped from an average of 15% at 3 days post-tamoxifen to an average of 3% at 7 weeks post-tamoxifen (Fig. 2B, *Black*). In contrast, the mTC progeny population, defined as tdTom^+^ cells not expressing Cox1, increased by ∼5-fold from an average of 4% of pancreatic ducts at 3 days post-tamoxifen to an average of nearly 17% at 7 weeks post-tamoxifen, reflecting the degree of mTC loss (Fig. 2B, *Pink*). These data support mTCs adopting a new identity in carcinomas.

### Transdifferentiation of mTCs to metaplastic neuroendocrine cells

To investigate the identity of the mTC progeny, we co-stained for tdTom together with markers of the known identities of other metaplastic ductal cells defined by *Schlesinger et al*. and *Benitz et al.*, including gastric pit-like (Gkn1^+^), gastric neck-like (Lectin GS-II^+^) ^29^, and metaplastic neuroendocrine cells (Syp^+^) ^7, 9, 17^.

The lack of co-expression of tdTom with Gkn1 or Lectin GS-II suggested that *KPF-P2f3T* tdTom^+^ mTC progeny had not taken on gastric pit-like cell or gastric neck-like cell identities (Fig. 2C and Supp. Fig. 2A). Instead, mTC progeny stained positive for Syp, revealing the acquisition of a metaplastic neuroendocrine cell (mNEC) identity (Fig. 2C, *Yellow arrows*). To determine if tdTom expression in cells with a neuroendocrine phenotype could be due to previously unnoticed recombination in pancreatic islets, we confirmed no co-localization of tdTom co-staining with Insulin (Ins) or in surrounding islet cells (Supp. Fig. 2B). Quantitation of the different time points for the *KPF-P2f3T* model confirmed a decrease of tdTom^+^/Cox1^+^ mTCs, (Fig. 2E, *Black*), with a concomitant ∼4-fold increase of tdTom^+^/Syp^+^ mNECs, (Fig. 2E, *Pink*) by 7 weeks post-tamoxifen (Fig. 2D, 2E). Interestingly, while there is a mNEC population in the metaplastic ducts in the *KF-P2f3T* neoplasia model, none expressed tdTom, suggesting that they did not derive from mTCs (Fig. 2D, 2E, *Green*) and had a distinct origin from those in the *KPF-P2f3T* mice.

### mNECs depend on the mTC population to develop in PDA but not PanIN models

Lineage tracing experiments show that mNECs in both neoplasia and carcinoma models are initially derived from acinar cells ^9^ (Supp. Fig 2C) but differ in that carcinoma mNECs derive from an mTC intermediate. As an independent test of these divergent origins, we took advantage of our *Pou2f3^CreERT/+^*driver being a knock-in to the *Pou2f3* allele, thus inactivating endogenous Pou2f3 activity ^20^. By creating a model homozygous for *Pou2f3^CreERT/CreERT^*, we effectively ablated *Pou2f3* (Supp. Fig. 3A, 3B), preventing mTC formation ^8^ and, therefore, any mTC progeny. Consistent with our previous observations, Cox1^+^ mTC and Syp^+^ mNEC populations were easily identifiable in the pancreatic lesions of our KF; Pou2f3^CreERT/+^ neoplasia model. Pou2f3 ablation in these mice eliminated the mTC population, as expected, while the mNEC population persisted (Fig. 3A), supporting their independent origins. In contrast, in the KPF; Pou2f3^CreERT/CreERT^ carcinoma mice, we found neither mTCs nor mNECs, confirming a shared dependency on Pou2f3 function (Fig. 3A, *Green arrows*).

**Figure 3.**
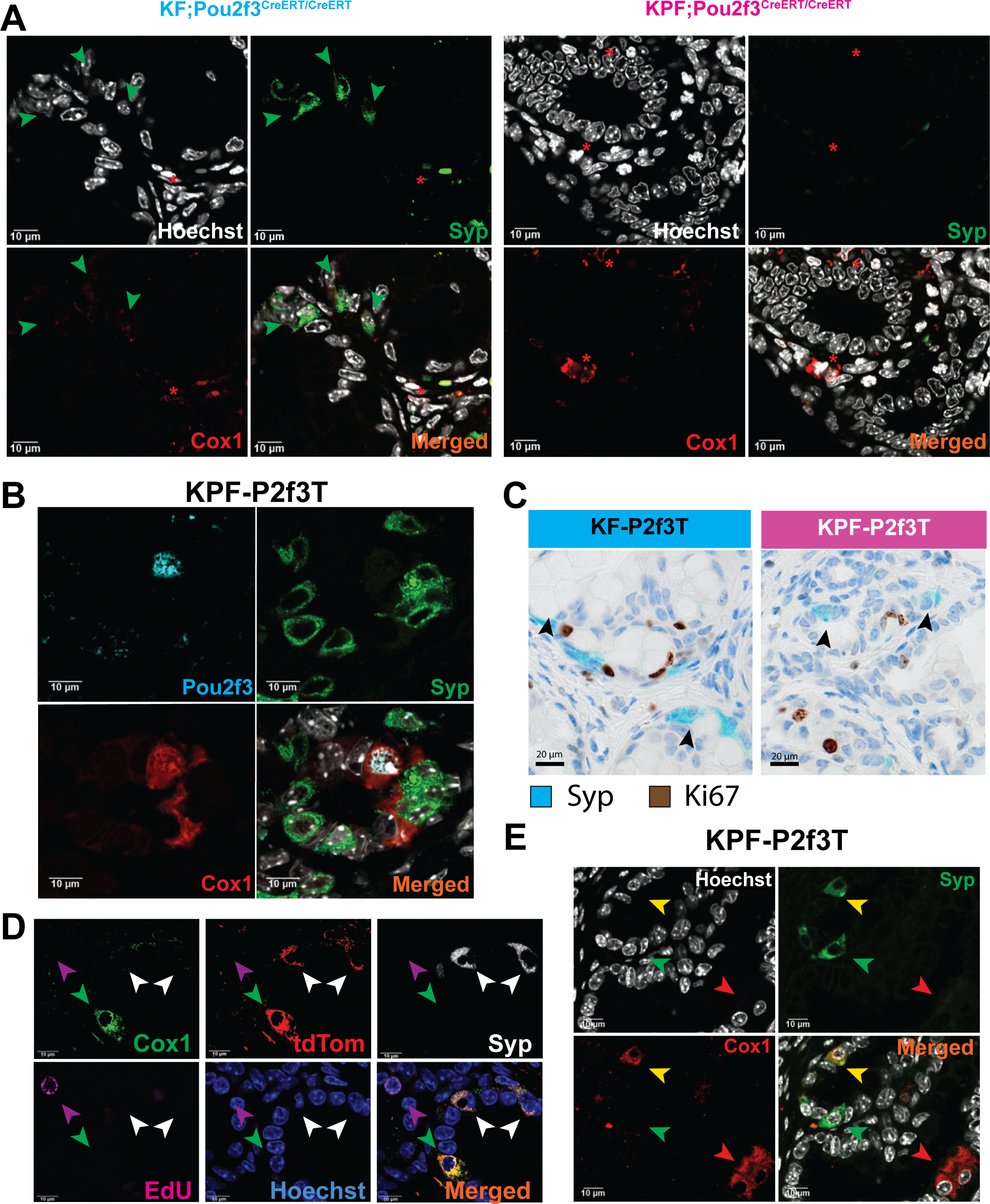
Characterizing the tuft cell to neuroendocrine cell transdifferentiation (TNT). IF staining of Cox1 (Red) and Syp (Green) in neoplasia and carcinoma models determines that mNECs in the *KPF-P2f3T* model relies on the mTC population. **A)** Cox1 (Red) and Syp (Green) markers are stained in Pou2f3 double knock-in models of pancreatic neoplasia and carcinoma: (Left) KF; Pou2f3^CreERT/CreERT^ and (Right) KPF; Pou2f3^CreERT/CreERT^, correspondingly to investigate the role of tuft cells in mNEC genesis. Green Arrows denote mNECs, Red Asterisks denote stromal Cox1^+^ staining identifying Cox1^+^ Immune cells. **B)** Pou2f3, the master regulator of tuft cells (Cyan) is stained with Cox1, marker of tuft cells (Red), and Syp, neuroendocrine marker (Green) to determine if mNECs also express Pou2f3. **C)** Dual IHC of Syp (Teal) and Ki67 (Brown) of neoplasia and carcinoma models of pancreas to determine proliferation of mNECs. **D)** Co-IF of mTC marker, Cox1 (Green), lineage tracer, tdTom (Red), mNEC marker, Syp (White), incorporated EdU (Magenta) to mark proliferating cells, and nuclear marker, Hoechst (Blue) in *KPF-P2f3T* mice treated with EdU over the course of 3 days post-tamoxifen treatment. **E)** Yellow arrows mark rare transitionary cells that are marked with both Syp (Green) and Cox1 (Red). Green arrow identifies cells that are only expressing Syp, mNECs. Red arrow marks cells only expressing Cox1, mTCs. Scale bars = 10 μm unless otherwise noted.

Lineage tracing and Pou2f3 ablation data in KPF mice support a model where mTCs give rise to mNECs. However, it was also possible that in the KPF model, both cell types express *Pou2f3* and require its activity. mNECs do not express *Pou2f3* in *Kras^G12D^*-driven neoplasia ^7, 22, 27^ but have not been examined in the KPF carcinoma model. To address this, we performed co-immunofluorescence for Pou2f3, together with Cox1 (mTCs) and Syp (mNECs). As expected, Pou2f3 was readily detectable in mTCs in both KF and KPF pancreata (Fig 1E, 3B). In contrast, we found no examples of Pou2f3^+^ mNECs in either system (Fig. 3B). The lack of identifiable recombination in early mNECs suggests that the increasing population of tdTom^+^ mNEC cells is dependent on the prior genesis of the mTC population.

### mTCs transdifferentiate into mNECs in PDA

The observation that mTCs act as a precursor cell to mNECs suggested two possible models for this relationship: mTCs act as a stem-like cell, giving rise to mNEC daughter cells, or alternatively, they directly transdifferentiate to mNECs. To distinguish these possibilities, we tested if mNEC genesis had undergone DNA replication, consistent with them arising from an asymmetric cell division of mTCs. mTCs rarely proliferate in neoplasia models ^30^, but their proliferative capacity has not been examined in a carcinoma model.

Initially, we co-stained for Ki67 and Syp and found that, like mTCs, mNECs were not in the cell cycle in *KF-P2f3T* or *KPF-P2f3T* mice (Fig. 3C, *Black arrows*). To test if a rare cell division event was necessary for mNEC genesis, we used a long-term EdU incorporation assay to determine if mNECs had gone through a prior cell division ^9, 11^. *KPF-P2f3T* mice were treated with tamoxifen once daily for 5 days at 11-12 weeks of age. We treated mice with EdU every 12 hours for 3 days and collected pancreata 12 hours after the last injection. Although we identified several EdU+ cells in the tissue (*Magenta arrows)*, neither Cox1^+^ mTCs (*Green arrows)* nor Syp^+^ TNTs (*White arrows*) had incorporated EdU (Fig. 3D), and, given the relatively long-time frame to allow for incorporation, were unlikely to have proliferated prior to their appearance.

To test the alternative that mTC-derived mNECs arise from the direct transdifferentiation of mTCs, we hypothesized that there would be cells that exist in mid-transition, co-expressing markers of both mTCs (Cox1) and mNECs (Syp). To identify transitionary cells, we co-stained Cox1 and Syp in *KPF-P2f3T* mice at 7 weeks post-tamoxifen treatment. Although rare, we identified cells expressing both Cox1 and Syp (Fig. 3E, *Yellow arrows*) exclusively in *KPF-P2f3T* mice. These transitionary cells were also EdU negative (Supp. Fig. 3C, *Yellow arrows*). Collectively, our data support a “Tuft-to-Neuroendocrine Transition” (TNT) transdifferentiation event, analogous to an epithelial-to-mesenchymal transition, exclusively in our pancreatic carcinoma model.

### Neoplastic mNECs have an enteroendocrine phenotype

While we have found mNECs in the pancreata of both neoplasia and carcinoma models, they appear to have distinct cellular origins. We confirmed that mNECs and mTCs in both models derive from acinar cells ^17, 22^ using an acinar cell lineage trace of *Pt1fa^Cre/+^; ROSA26^LSL-YFP/+^* in neoplasia and carcinoma models. We show co-localization with GFP (to mark acinar cells and progeny) and either Cox1 (mTCs) or Syp (mNECs) for both models (Supp. Fig. 2C). Although mNECs derive from acinar cells in neoplasia and carcinoma, in the *KPF-P2f3T* carcinoma model, mNECs require the genesis of an mTC intermediate. This unexpected observation led us to ask if mNECs in the *KF-P2f3T* neoplasia model and the *KPF-P2f3T* carcinoma model are distinct identities.

Two types of mNECs are described in human pancreatic disease: neuroendocrine-like cells ^21^, also designated enteroendocrine cells (EECs) ^17, 22^, marked by their expression of islet cell hormones, and neural-like progenitor cells (NRPs) ^21^, also called ductal neuroendocrine cells ^7, 23^, marked by the expression of genes associated with neural development. EECs are distinct from tuft cells in their origins in pancreatic disease ^17, 22^ in that they do not rely on Pou2f3 activity ^17, 22, 31^, and express markers of normal islet cells ^21, 22^. To determine if mNECs in our models bore an EEC phenotype, we examined islet cell hormone expression ^21^ by co-IF of wide-spectrum cytokeratin (PanCK) and Syp together with either glucagon (Gcg), insulin (Ins), somatostatin (Sst), pancreatic polypeptide (Ppy), and ghrelin (Ghrl), markers of α, β, δ, γ, and ε islet cells, respectively ^32^.

Since the KF mNECs do not express a lineage tracer, we confined our analysis to PanCK^+^/Syp^+^ cells to distinguish them from singular islet cells that can become trapped within the diseased tissue (Figs. 4A-E & Supp. Fig. 3D). In the *KF-P2f3T* neoplasia model, we consistently found isolated mTC-independent mNECs expressing islet cell identity markers apart from the β cell markers Ins and serotonin (5-HT) (Fig. 4B and Supp. Fig. 2B). In contrast, we found no islet hormone expression in the mTC-derived mNECs in the *KPF-P2f3T* mouse. These data indicated that mNECs in our KF-P2f3T neoplasia model, but not in our *KPF-P2f3T* carcinoma model, are likely EECs, similar to those found in other models of pancreatic disease ^17, 21, 22^.

**Figure 4.**
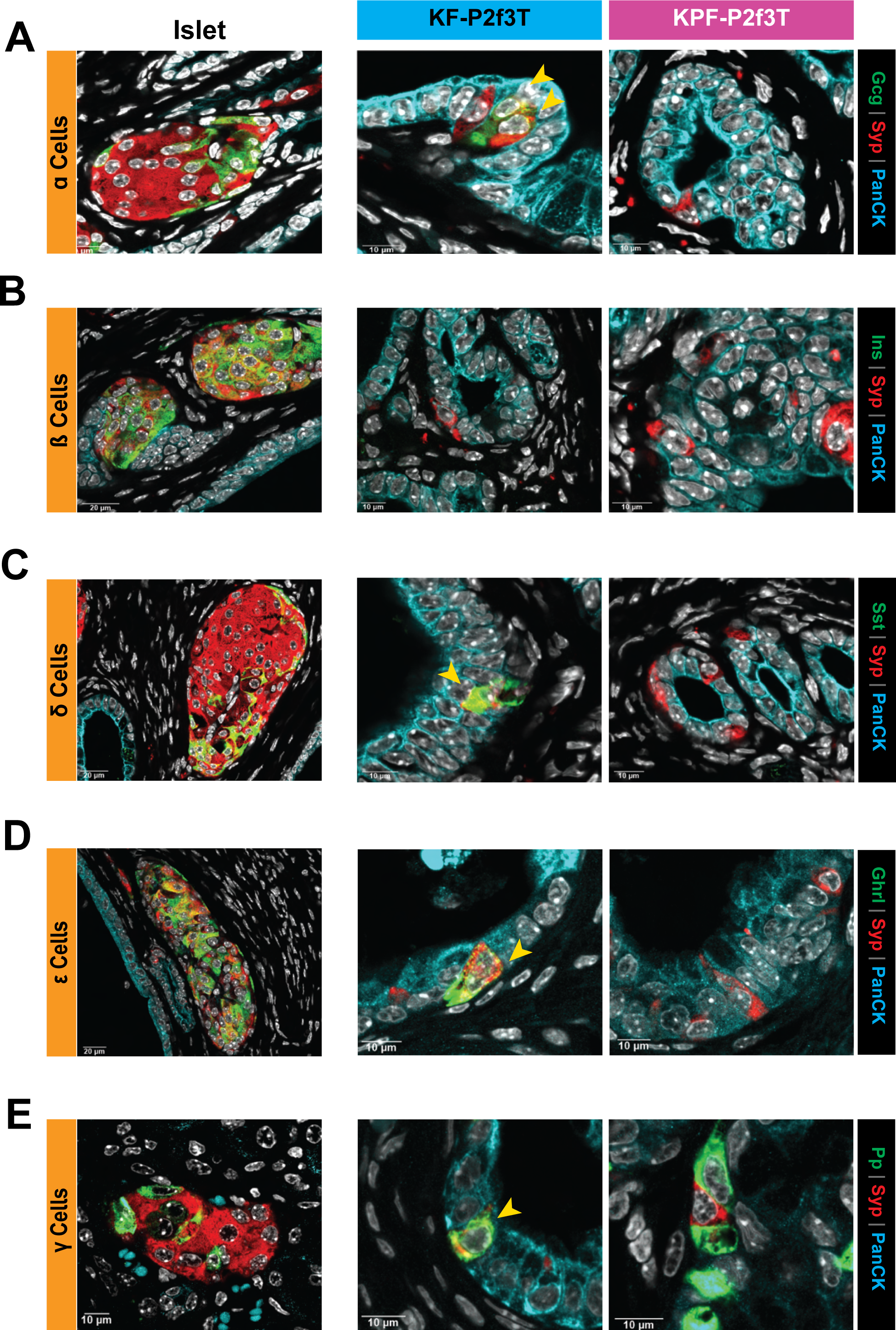
Identifying distinctions between mNECs in the progression of PDA. Neoplasia and carcinoma PDA models were stained with markers of different islet compartments to determine differences between mNECs between the *KF-P2f3T* and *KPF-P2f3T* models. **B)** Islets and lesions from *KF-P2f3T* and *KPF-P2f3T* are shown to determine co-localization of Syp (Red) and markers of the different compartments of the Islet (Green). PanCK (Cyan) was used to distinguish islet cells from mNECs, and Hoechst was used to mark the nucleus of each cell. Glucagon (Gcg), Insulin (Ins), Somatostatin (Sst), Ghrelin (Ghrl), and Pancreatic Polypeptide (Pp) were stained to identify the α cells, β cells, δ cells, ε cells and γ cells respectively. Scale bars = 10 μm unless otherwise noted.

### Cells derived from TNT persist in pancreatic carcinoma

Having established that mNECs in KF neoplasia are likely EECs, whereas those in the KPF carcinoma are not, we hypothesized that the TNT-derived mNECs are neural-like progenitor cells (NRPs), as identified in human PDA ^21^. In human PDA, NRPs correlate with poor survival and drug resistance ^23^. Initially, we asked if our TNT mNECs persist in frank carcinoma, as NRPs do in patients. To test this, we extended the trace time post-tamoxifen treatment to 12 weeks and examined tdTom expression in areas of undifferentiated carcinoma, confirmed by histology (data not shown). As expected, tuft cells were absent in regions of carcinoma, as indicated by IHC for Dclk1 (Fig. 5A, *Left*). However, when we stained adjacent sections for tdTom, we easily found mTC progeny cells scattered throughout the tumors (Fig. 5A, *Right*). Using co-IF, we confirmed that the identity of these mTC progeny were indeed Syp-expressing mNECs (Fig. 5B). We conclude that not only are mTCs undergoing TNT, but the mTC-derived mNECs persist within dedifferentiated carcinoma, consistent with NRPs in PDA patients ^21^.

**Figure 5.**
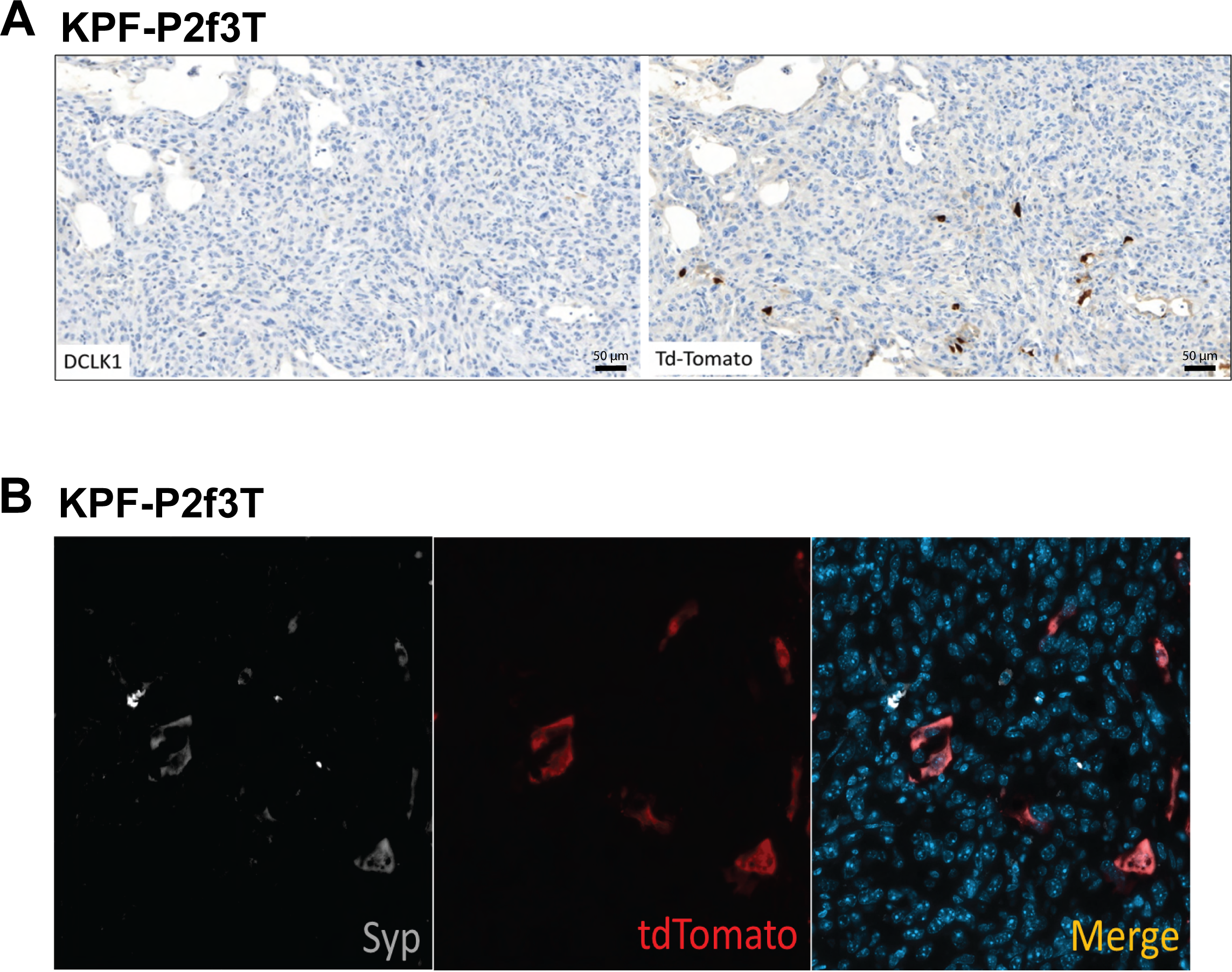
mTC-derived mNECs are identified in PDA Carcinoma. IHC and IF are used to confirm the identity of mTC Progeny in dedifferentiated carcinoma of *KPF-P2f3T* model. **A)** IHC of dedifferentiated carcinoma stained with Dclk1 (*Left*) and tdTomato (*Right*) in Brown. **B)** Dedifferentiated carcinoma of *KPF-P2f3T* model is stained with Hoechst (Cyan), Syp (White), and tdTomato (Red) to determine identity of mTC progeny. Scale bars = 50 μm

### TNT gives rise to Nrxn3^+^ neural-like progenitor cells in pancreatic carcinoma

Neurexin 3 (Nrxn3) is a marker of neural-like progenitor cells (NRPs) linked with poor prognosis and chemotherapy resistance in multiple cancers, including colon, lung, and pancreatic cancer ^21, 33, 34^. The persistence of mNECs in carcinoma led us to examine the expression of Nrxn3 in the TNT-derived mNECs in the KPF carcinoma model. KF neoplasia and KPF carcinoma tissues were co-stained for Nrxn3, Syp, and PanCK to distinguish mNECs with an NRP identity. As expected, *KF-P2f3T* EECs did not express Nrxn3 (Fig. 6A and 6B, *Top*). However, in the *KPF-P2f3T* carcinoma model, Nrxn3, Syp, and PanCK co-localization confirmed that TNT-derived mNECs cells were indeed NRPs both in areas of neoplasia (Fig. 6A and 6B, *Bottom*) and carcinoma (Fig. 6C).

**Figure 6.**
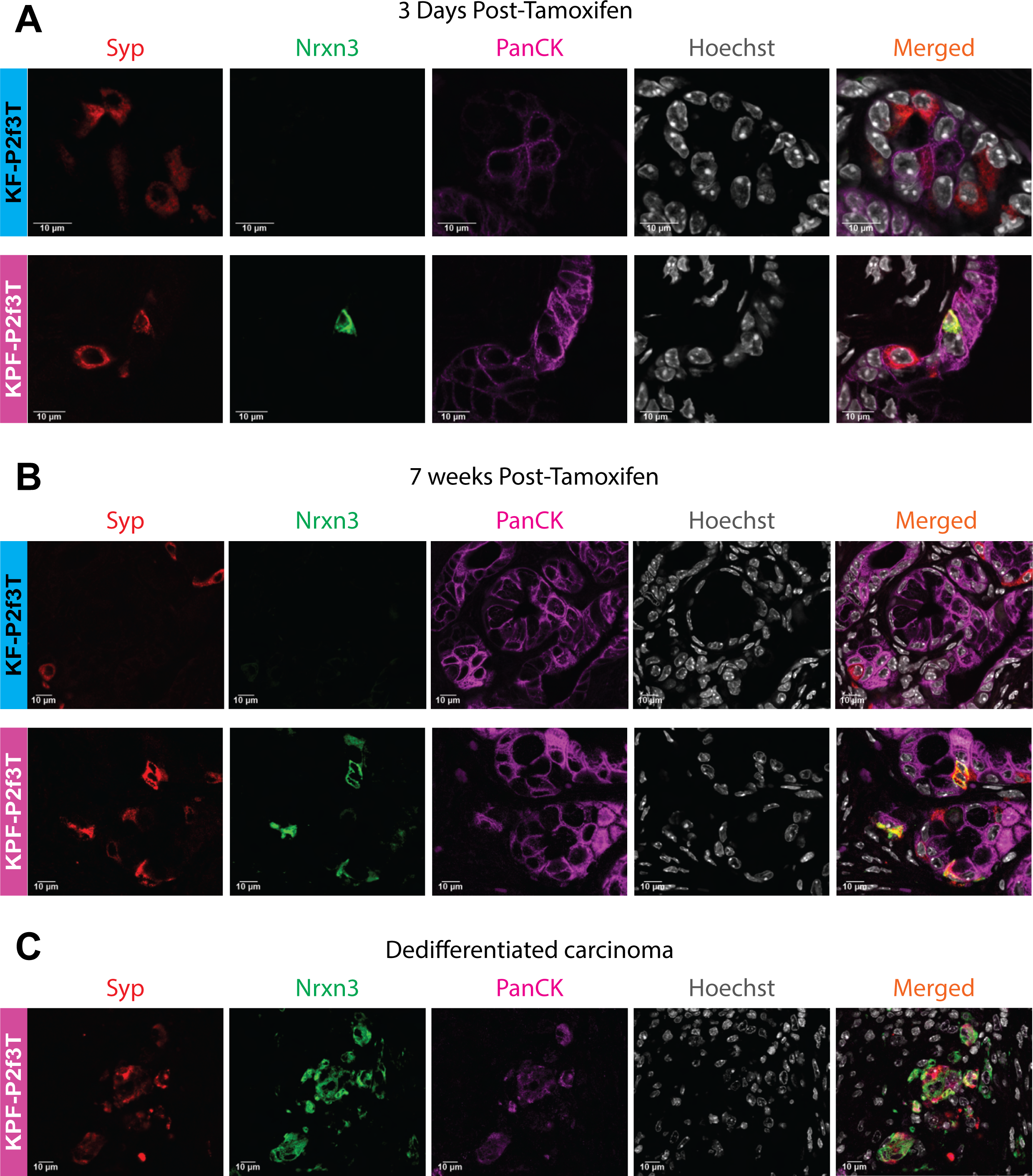
Identification of mTC-derived Nrxn3+ NRPs in Carcinoma. Co-staining of Syp (Red), Nrxn3 (Green), PanCK (Magenta), and Hoechst (White) in *KF-P2f3T* neoplasia model and *KPF-P2f3T* carcinoma model at **A)** 3 days post-tamoxifen, **B)** 7 weeks post-tamoxifen, and **C)** *KPF-P2f3T* dedifferentiated carcinoma. Scale bars = 10 μm unless otherwise noted.

### The Tuft to Neuroendocrine transition is dependent upon *Myc* activity

Pancreatic parenchyma-wide *Myc* activity induces the formation of mNECs in a KC model of neoplasia ^23^, similar to the prostate and lung ^35–38^. NRPs in patient samples are marked by high levels of MYC activity in cancers of the pancreas and other tissues ^21, 23, 35, 36^. We hypothesized that targeting *Myc* activity specifically to mTCs in the *KF-P2f3T* neoplasia mouse, where TNT is not otherwise observed, would be sufficient to induce NRP genesis. To accomplish this, we crossed the *Myc* Cre-dependent overexpression mouse line, ROSA26^LSL- cMyc/+^, into the *KF-P2f3T* model (designated KF-MOE) (Supp. Fig. 4A). We treated these mice with cerulein to induce transformation and mTC genesis, then treated with tamoxifen 3 weeks later to activate the tdTom reporter and induce upregulation of *Myc* specifically in the mTCs. As previously observed, the control *KF-P2f3T* mice had no mTC-derived mNECs. In contrast, we readily found mTC-derived mNEC genesis in KF-MOE tissue as early as 3 days post-induction (Fig. 7A and Supp. Fig. 4C), *Yellow arrows*), with a 6-fold increase in the tdTom^+^ mNEC population 7 weeks post-tamoxifen treatment (Fig 7C, *Pink*). Finally, to determine if the mNECs in the *KF-MOE* mice were NRPs, we co-stained tissue 7 weeks post-tamoxifen for Nrxn3, Syp, and tdTom (Fig. 7B). Indeed, we consistently found NRP identity in the neuroendocrine population induced by overexpressing *Myc* in mTCs. Thus, *Myc* expression targeted to mTCs is sufficient to induce transdifferentiation to NRPs in a neoplasia model where NRPs are not observed otherwise.

**Figure 7.**
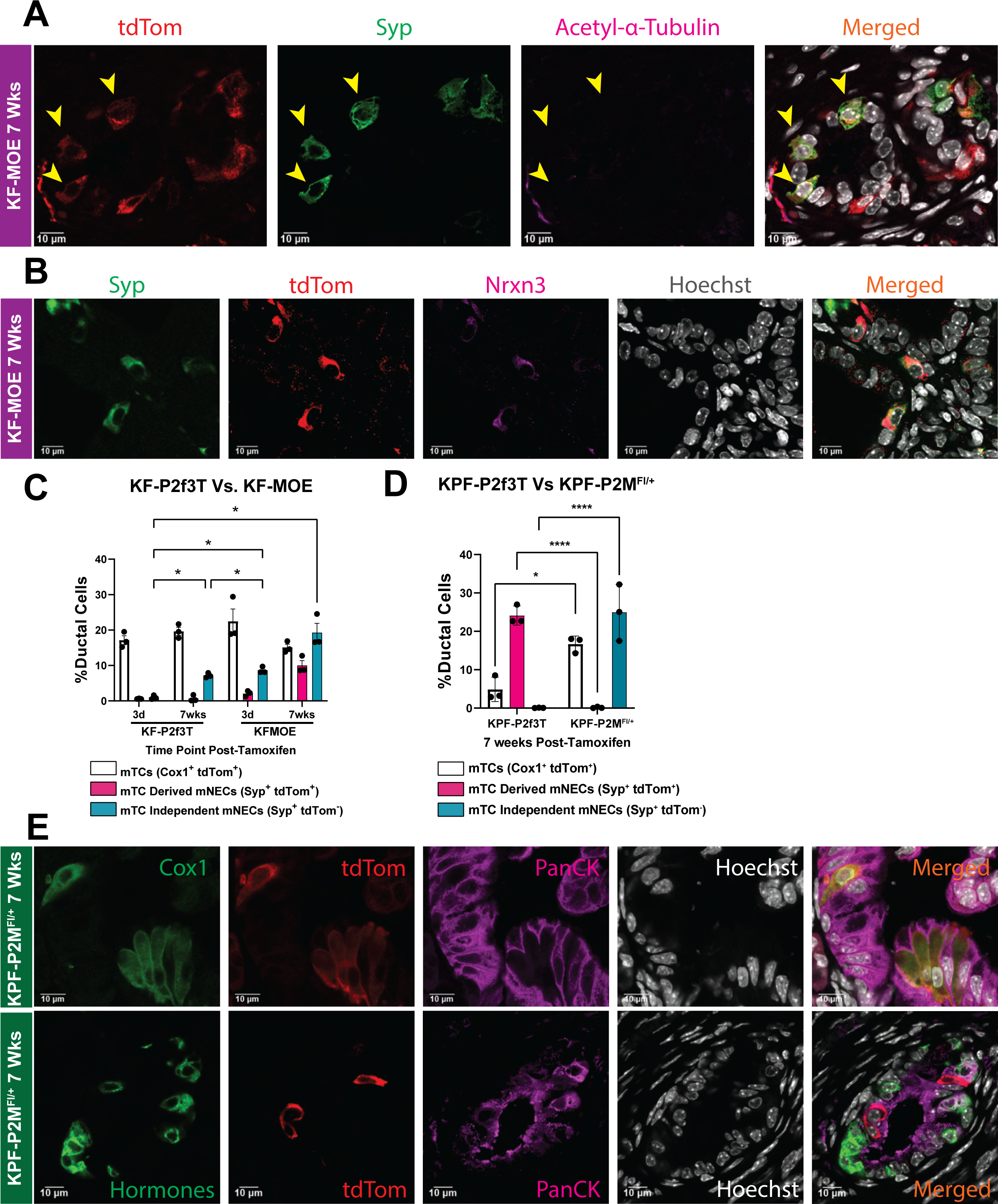
cMyc Mediated development of mNECs. Overexpression and knockdown of *Myc* in neoplasia and carcinoma models show dependence of *Myc* expression on transition of mTCs to NRPs in PDA progression. **A)** *KF-MOE* 7 weeks post-tamoxifen treatment are stained with tdTom to identify cells recombined by tamoxifen treatment and cells overexpressing *Myc* (Red), Syp to mark neuroendocrine cells (Green), Acetylated-α-tubulin to mark filamentous actin of cells (Magenta), and Hoechst to identify nucleus of all cells (White) to detect mTC-derived mNECs. **B)** KF-MOE 7 weeks post-tamoxifen treatment tissue is stained for co-IF to determine expression of Syp (Green), tdTom (Red), Nrxn3 (Magenta), and Hoechst (White). **C)** Quantification of mTCs (Cox1+ tdTom+) in White, mTC derived mNECs (Syp+ tdTom+) in Pink and mTC independent mNECs (Syp+ tdTom-) in Green for *KF-P2f3T* and *KF-MOE* models. Mice were sac’d after 3 days and 7 weeks post tamoxifen treatment to label tuft cells with tdTomato and overexpress cMyc specifically in tuft cells. (n = 3/group) p-values were calculated using a two-way ANOVA on Prism GraphPad 7. * = p < 0.05. D) Quantification of mTCs (White), mTC Derived mNECs (Pink), and mTC Independent, (Green) in *KPF-P2f3T* and *KPF-P2M^Fl/+^* models. Mice were sac’d at 7 weeks post tamoxifen treatment to label tuft cells with tdTom and flox *cMyc* expression in tuft cells specifically. (n = 3/group) p-values were calculated using a two-way ANOVA on Prism GraphPad 7. * = p < 0.05, **= p<0.005. **E)** Co-staining of Cox1 (Green), tdTom (Red), PanCK (Magenta), and Hoechst (White) in *KPF-P2M^Fl/+^* at 7 weeks post-tamoxifen treatment to identify co-localization of mTC marker, Cox1, with tdTom (*Top*). Co-staining of combined antibodies for Ghrl, Gcg, Sst, Ppy, and Ins to mark cells expressing “Hormones” to identify EECs (Green), tdTom used to identify cells recombined and *Myc* knockout cells (Red), PanCK to mark pancreatic epithelium/lesions (Magenta), and Hoechst to mark nucleus of cells (White) in *KPF-P2M^Fl/+^* at 7 weeks post-tamoxifen treatment to identify EECs and tdTom+ cells as separate cell populations when *Myc* is conditionally knocked out in carcinoma model. Scale bars = 10 μm unless otherwise noted.

Having shown that Myc activity is sufficient to drive TNT, we tested if it was necessary for the process. To accomplish this, we crossed a conditional *Myc* knock-out allele (*Myc*^fl/+^) into the *KPF-P2f3T* model (designated *KPF-P2M*^fl/+^) (Supp. Fig. 4B). These mice were treated with tamoxifen to simultaneously activate the tdTom reporter and reduce *Myc* activity specifically in the mTCs. As previously, the *KPF-P2f3T* control at 7 weeks post-tamoxifen showed a lack of mTCs and an increased population of mTC-derived mNECs (Fig. 7D). In contrast, in the *KPF-P2M*^fl/+^ pancreata, we could easily identify Cox1^+^/tdTom^+^ mTCs at 7 weeks post-tamoxifen (Fig. 7D&E) and a lack of Syp^+^/tdTom^+^ mTC-derived mNECs.

Unexpectedly, these mice had a significant population of Syp^+^/tdTom^-^ mTC-independent mNECs (Fig. 7D; Supp. Fig 4D). To test if the mTC-independent mNECs were EECs, we combined antibodies for Ghrl, Gcg, Sst, Ppy, and Ins to identify any hormone-positive cells simultaneously and co-stained for tdTom (Fig. 7E). Indeed, the tdTom^-^ mNECs in the *KPF-P2M*^fl/+^ model expressed islet hormones, confirming their identity as EECs. We conclude that *Myc* activity is necessary and sufficient to induce mTC transdifferentiation into NRPs.

## Discussion

In the pancreas, prolonged injury or expression of oncogenic *Kras* leads to acinar-to-ductal metaplasia, with metaplastic ducts comprised of at least 6 different cell types: gastric pit-like cells, gastric chief-like cells, gastric SPEM/neck-like cells ^6^, senescent duct cells, metaplastic neuroendocrine cells (mNECs), and metaplastic tuft cells (mTCs) ^7, 9, 17^. The functions of these different metaplastic cell types have only begun to be recognized. For instance, normal tuft cells in other organs combat infection ^14^ and promote wound healing ^11, 30, 39^. Consistent with their overall role in maintaining normal tissue homeostasis, mTCs suppress the progression of pancreatic neoplasia to adenocarcinoma ^8, 10^. In contrast, mNECs in the pancreas are associated with tumor cell-neuron cross talk and promote tumor cell proliferation ^40^.

While mTCs functionally hinder the progression of neoplasia to pancreatic cancer, they also have been theorized to be tumor progenitor cells in the pancreas^11, 30, 41^, intestine ^42^, and stomach ^43^. Doublecortin-like kinase (Dclk1), a common tuft cell marker, is also a marker of progenitor cells in the pancreas and intestine ^11, 30, 42^. Dclk1 is a microtubule kinase highly expressed in neurons and is linked with neuronal survival and migration in other organs ^44^.

Besides Dclk1, mTCs express other stem cell markers such as CD133, CD24, and CD44 ^11^, which, together with their low proliferation rate ^11, 30, 42, 45^, support their role as putative cancer stem cells. Consistent with a facultative progenitor cell function, depleting mTCs delays tissue repair after experimental pancreatitis ^30, 42^. However, contrary to their proposed role as cancer-initiating cells in PDA, mTC numbers diminish as neoplasia progresses to carcinoma ^9, 11^, inconsistent with their mandatory role in the genesis of new tumors.

We theorized many possible reasons why mTCs disappear through progression. With little evidence that they are dying or being outcompeted by more proliferative cell types, we focused on the possibility that they were changing their cellular identity. Using a unique dual recombinase mouse model to trace mTC fate through progression, we found that mTCs were stable in *Kras^G12D^*-driven neoplasia but transdifferentiated to a mNEC identity in a *Kras^G12D^*;

*Tp53^+/-^* carcinoma model. Importantly, we cannot distinguish whether it is progression to a less differentiated adenocarcinoma in general or the deletion of a *Tp53* allele specifically that is responsible for driving this transition. However, given that we do not observe mTCs or their progeny in carcinomas derived from *Kras^G12D^* expression only (data not shown), this suggests an association with general tumor progression, agnostic to specific tumor suppressor gene mutations.

Pancreatic tumor cells expressing neural/neuroendocrine markers in human PDA and are comprised of two different subsets: neuroendocrine-like or enteroendocrine cells (EECs) ^7, 22^ and neural-like progenitor cells (NRPs) ^21^, the latter of which is associated with poor outcomes. EECs express hormone markers typically confined to islet cells ^17, 21, 22^, whereas NRPs express markers of neural progenitor cells, including Nrxn3 ^21^. We found that mNECs in *Kras^G12D^*-driven neoplasia arise independent of mTCs and take on an islet hormone-positive EEC identity, consistent with previous data showing EECs in metaplasia are independent of mTCs and do not depend on *Pou2f3* to develop ^22^. In contrast, mNECs in *Kras^G12D^; Trp53^+/-^* carcinoma do derive from mTCs and have an Nrxn3^+^ NRP phenotype.

EECs share a seemingly similar fate as mTCs in that they are present at early stages of ADM but diminish as PDA progresses to dedifferentiated carcinoma ^22^. Our data support an association of EECs with early oncogenesis as they are largely confined to our neoplasia model and absent in our carcinoma model. Unexpectedly, ablation of a single allele of *Myc* in the carcinoma model mTCs allowed EECs to reemerge, suggesting an inverse relationship between EECs and NRPs.

Mouse mNECs (both EECs and NRPs) derive from acinar cells ^17, 22, 23^, but NRPs require an mTC intermediate. NRPs are associated with high *Myc* activity in PDA and other carcinomas such as lung, colon, and prostate cancer ^21, 23, 35, 36^. Supporting the orthologous nature of mouse NRP cells, upregulating *Myc* expression specifically in mTCs in our neoplasia model was indeed sufficient to drive NRP transdifferentiation whereas reducing *Myc* activity in the mTCs of our carcinoma model inhibited the process. Together, these data suggest that *Myc* is both necessary and sufficient for a “tuft to neural progenitor transition” (TNT) in PDA. NRPs contribute to tumorigenicity ^46^ and advanced carcinomas frequently express neural characteristics ^47^, consistent with their association with poor outcomes in PDA patients ^21, 23^ and our observation that NRPs are long-lived, persisting within poorly differentiated carcinomas. Even so, we have not yet observed conditions under which murine NRPs proliferate so as to constitute a significant population of carcinoma cells. It is possible that NRPs being resistant to chemotherapy may be selected for confronted with treatment ^21, 23^, and possibly proliferate subsequently in that environment.

The coexistence of tuft cells and neuroendocrine cellular subpopulations is a characteristic of several carcinoma types. Small cell lung cancer (SCLC) can be classified into subtypes that include a tuft cell-like subtype based on POU2F3 expression and two neuroendocrine subtypes based on ASCL1 and NEUROD1 expression ^26^. Tuft cell-like SCLC transitions to neuroendocrine subtypes as SCLC tumors lose*Tp53* and have an increased activity of *Myc* ^37^. In addition to this neural transition, these cells are characterized by the gain of EMT markers in SCLC ^35, 37^. Neuroendocrine prostate cancer (NEPC) NRPs are also driven by *Myc* expression and *Tp53* deletion ^38, 48^. In the setting of prostate cancer, NEPC is derived from epithelial cells^36^ in prostate adenocarcinoma after the treatment of hormone therapy and is associated with worse outcomes ^38^.

In summary, our study reveals a novel type of transdifferentiation of metaplastic tuft cells to neural-like progenitor cells, shifting their role from suppressing tumor progression through their chemosensory function in early-stage disease, to being associated with poor outcomes in the later stages after TNT. We found that the TNT requires *Myc* activity ^49–51^ and possibly the loss of *Tp53* function^37^, associations that have been observed in other tumor types ^27, 35–37^. While this does not settle the debate as to whether mTCs can act as tumor progenitor cells, the NRP progeny of mTCs can persist long term as part of a poorly differentiated carcinoma, though their lack of proliferation under unperturbed conditions makes it unlikely that they contribute significantly to overall tumor burden. The observation that NRPs are resistant to chemotherapy and their numbers are associated with worse outcome and tumor recurrence^21, 23^ suggest that, when challenged, their role may shift to a more traditional tumor initiating cell. Other studies suggest that mNECs instead play a supportive role for other carcinoma cells, promoting their expansion and possibly contributing to their survival under stress ^40^, though whether these are functions common to both EECs and NRPs has not been determined. While the specific functions of NRPs in PDA remain under investigation, understanding the mechanisms controlling their genesis suggests that one possible way to target these aggressive carcinoma cells is to reverse the TNT process to restore their mTC tumor suppressing role.

## Methods

### Mouse Models

#### mTC Lineage tracing

*Pou2f3^CreERT/+^* ^19^, *ROSA26^LSL-TdTomato/+ 28^*. These were combined to generate *Ptf1a^FlpO/+^; FSF-KRAS^G12D/+ 18, 52^; Pou2f3^CreERT/+ 19^; ROSA26^LSL-TdTomato/+^ (KF-P2f3T) and Ptf1a^FlpO/+^; FSF-KRAS^G12D/+^; Trp53^Frt-Exons 2 to 5-Frt/+ 18, 52^; Pou2f3^CreERT/+ 19^; ROSA26^LSL-TdTomato/+^* (*KPF-P2f3T*) mice. We induced pancreatic damage in the 8- to 9-week-old *KF-P2f3T* mice with a once-daily intraperitoneal injection of cerulein (46-1-50; American Peptide Company, Inc, Sunnyvale, CA) at a dose of 250 μg/kg for a total of five days. To induce recombination of ROSA26^LSL-TdTomato/+^ in *KF-P2f3T* and *KPF-P2f3T* models, 11- to 12-week-old mice were treated with 5mg of Tamoxifen (T5648; Millipore-Sigma, St Louis, MO) via oral gavage, once a day, for 5 days. Tamoxifen was dissolved in corn oil before treating mice. Pancreata were then harvested 1-day post-tamoxifen, three days post-tamoxifen, and seven weeks post-tamoxifen to capture different stages of PDA in mice.

#### Acinar-derived mTCs and mNECs

*Ptf1a^Cre+^; LSL-KRAS^G12D/+^; LSL-Trp53^R172H/+^; ROSA26^LSL-YFP/+^* ^9^ were combined to generate *Ptf1a^Cre+^; LSL-KRAS^G12D/+^, ROSA26^LSL-YFP/+^ (*KCY) and *Ptf1a^Cre+^; LSL-KRAS^G12D/+^; LSL-Trp53^R172H/+^; ROSA26^LSL-YFP/+^* (KPCY) mice to allow lineage trace of epithelial compartment. KCY mice were treated with Cerulein at 9 weeks old and allowed to recover for 3 weeks. KCY and KPCY mice were treated with Tamoxifen to induce YFP expression in Ptf1a+ cells at twelve weeks of age.

#### mTC Ablation

A homozygous CreERT knock-in inserted into the *Pou2f3* locus was used to induce metaplastic tuft cell ablation in KF and KPF mice. 8- to 9-week-old *Ptf1a^FlpO/+^; FSF-KRAS^G12D/+ 18, 52^; Pou2f3^CreERT/CreERT^* (KF; Pou2f3^CreERT/CreERT^) mice were treated with cerulein via intraperitoneal injection at a dose of 250 μg/kg for a total of 5 days to induce pancreatic damage. *Ptf1a*^FlpO^*^/+^; FSF-KRAS^G12D/+^; Trp53^Frt-Exons 2 to 5-Frt/+^; ^18, 52^ Pou2f3^CreERT/CreERT^* (KPF; Pou2f3^CreERT/CreERT^) did not require cerulein to cause damage. Pancreata were harvested at 12 weeks of age to determine ablation of mTCs in both models.

#### cMyc overexpression in mTCs

To Investigate the role of *cMyc* in the transdifferentiation of mTCs into mNECs, we bred mouse lines to create the genotype: *Ptf1a^FlpO/+^, KRAS^FSF-G12D/+^, Pou2f3^CreERT/+ 16, 20^*, *ROSA26^LSL-TdTomato/+^, ROSA26 ^LSL-cMyc/+ 23^*. These were combined to generate KF-MOE mice comprised of *Ptf1a^FlpO/+^; FSF-KRAS^G12D/+ 18, 52^; Pou2f3^CreERT/+ 20^; ROSA26^LSL-TdTomato/+^; ROSA26 ^LSL-cMyc/+ 23^*. KF-MOE mice overexpressed *cMyc* and tdTomato reporter specifically in mTCs in the KF neoplasia model. Pancreatic damage was induced in 8- to 9-week-old KF-MOE mice with a once-daily intraperitoneal injection of cerulein (46-1-50; American Peptide Company, Inc, Sunnyvale, CA) at a dose of 250 μg/kg for a total of 5 days. To induce recombination of *ROSA26^LSL-TdTomato/+^*and *ROSA26^LSL-cMyc/+ 23^* in the KF-MOE model, eleven- to twelve-week-old mice were treated with 5mg of Tamoxifen (T5648; Millipore-Sigma, St Louis, MO) via oral gavage, once a day, for 5 days. Tamoxifen was dissolved in corn oil before treating mice. We harvested pancreata 3 days post-tamoxifen and seven weeks post-tamoxifen to investigate the role of *cMyc* in mTC-derived mNECs.

#### cMyc flox in mTCs

To Investigate the role of *cMyc* in the transdifferentiation of mTCs into mNECs, we bred mouse lines to create the genotype: *Ptf1a^FlpO/+^, KRAS^FSF-G12D/+^, Trp53^Frt-Exons 2 to 5-Frt/+ 18, 52^, Pou2f3^CreERT/+ 16, 20^*, *ROSA26^LSL-TdTomato/+^, Myc^Flox/+ 23^*. These were combined to generate *KPF-P2M^Fl/+^*mice comprised of *Ptf1a^FlpO/+^; FSF-KRAS^G12D/+ 18, 52^; Trp53^Frt-Exons 2 to 5-Frt/+^ Pou2f3^CreERT/+ 20^; ROSA26^LSL-TdTomato/+^; Myc^Flox/+ 23^*. *KPF-P2M^Fl/+^* mice reduced *Myc* activity and expressed the tdTomato reporter specifically in mTCs in the KPF PDA carcinoma model. To induce recombination of *ROSA26^LSL-TdTomato/+^*and *Myc^Flox/+ 23^* in the *KPF-P2M^Fl/+^* model, eleven- to twelve-week-old mice were treated with 5mg of Tamoxifen (T5648; Millipore-Sigma, St Louis, MO) via oral gavage, once a day, for 5 days. Tamoxifen was dissolved in corn oil before treating mice. We harvested pancreata seven weeks post-tamoxifen to investigate the role of *Myc* in mTC-derived mNECs.

### Immunohistochemistry (IHC)

Mouse pancreas tissues were fixed overnight in a Z-fix solution (NC9050753; Anatech Ltd, Battle Creek, MI). The pancreas tissues were processed using a Leica ASP300S tissue processor (Buffalo Grove, IL). Sections (4 μm) of paraffin-embedded tissue were stained for target proteins (Table 1) of interest using the Discovery Ultra XT auto Stainer (Ventana Medical Systems Inc, Tucson, AZ). IHC slides were imaged and stitched together by a 3DHisTech scanner (Perkin Elmer, Seattle, WA) using a ×20 objective lens.

### Immunofluorescence (IF)

Tissues were prepared as Z-fixed paraffin-embedded blocks, as stated above. In increased dilutions, sectioned tissue was deparaffinized using xylene and a series of ethanol. We boiled the slides in a pH 6.0 Citrate buffer solution for 5 minutes. Slides were allowed to cool at room temperature for 20 mins, then washed in 0.1% tritonX + PBS solution for 15 mins three times for a total of 45 mins. The sections were then blocked in a donkey block solution of 5% donkey serum/1% bovine serum albumin (BSA) in PBS for 1 hour. Following the block, sections were incubated overnight with primary antibodies, as stated in **Supp. Table 1**. The following day, we washed in a 0.1% TritonX + PBS solution for 15 mins thrice. Then Alexa Fluor-conjugated secondary antibodies (1:500, Invitrogen, Carlsbad, CA) were added and incubated at RT for 1hr in the dark. Slides were then washed once more with 0.1% TritonX + PBS + Hoechst 33342 (62249, 1:10000, Thermo Scientific, Carlsbad, CA) solution for 15 mins thrice to remove excess secondary and label nuclei. Lastly, slides were briefly rinsed in deionized water, mounted with Prolong Diamond antifade mountant (P36961; Fisher), and dried overnight at room temperature before imaging.

### EdU Proliferation Assay

The “Click-iT™ Plus EdU Cell Proliferation Kit for Imaging, Alexa Fluor™ 488 dye” (ThermoFisher, Catalog number: C10637) was used to label cells that have undergone proliferation for a specific time period ^53^. We dissolved EdU in PBS to a concentration of 1mg/mL. *KPF-P2f3T* mice were treated with tamoxifen at 11- to 12-weeks of age once daily for 5 days. Mice were treated with 100uL of EdU injected via I.P. injection every 12 hours for 3 days post tamoxifen labeling. We harvested the pancreata from each mouse the following morning. We fixed and stained each pancreas as mentioned in the IF protocol above just until after secondary antibodies were added and washed in 0.1% TritonX + PBS + Hoechst 33342 (62249, 1:10000, Thermo Scientific, Carlsbad, CA) solution for 15 mins three times to removed excess secondary and label nuclei. We detected EdU by making a cocktail using the following components provided by the kit: 440uL of 1x Click-iT Reaction buffer, 10uL of Copper Protectant, 1.2uL of Alexa Fluor Picolyl, and 50uL of Reaction buffer additive.

Reagents must be added in the order listed. 100uL of the cocktail was added to each slide and incubated for 30 minutes before washing with 3% BSA in PBS. We rinsed the slides with ddH2O, mounted them with Prolong Diamond antifade mountant (P36961; Fisher), and dried them overnight at room temperature before imaging.

### Quantitation

HALO Software (Indica Labs, Corrales, NM) was used to quantify whole tissue scanned sections. Quantification excluded blood vessels, lymph nodes, and adipose/connective tissue. We took multiple sections from each tissue throughout the block and averaged them to produce a data point. Through this software, we can train the algorithm to recognize specific areas of tissue and allow us to determine marker expression in different regions of the pancreas using the whole section. We did image analysis as per Indica Labs’ recommendations.

## Acknowledgements

The authors would like to thank Marina Pasca di Magliano for helpful discussions. This work was supported by R01CA235141, U01CA224145 and U01CA274154 (to HCC). DJSE was supported by the University of Michigan’s Rackham Merit Fellowship, NCI F31CA271576, and NCI T32CA009676. KED is supported by the Vanderbilt Ingram Cancer Center Support Grant (NIH/NCI P30CA068485), the Vanderbilt-Ingram Cancer Center SPORE in Gastrointestinal Cancer (NIH/NCI P50CA236733), the Vanderbilt Digestive Disease Research Center (NIH/NIDDK P30DK058404), an American Gastroenterological Association Research Scholar Award (AGA2021-13-02), NIH/NIGMS R35GM142709, The Department of Defense (DOD W81XWH2211121), The Sky Foundation, Inc (AWD00000079), and Linda’s Hope (Nashville, TN). RCS is supported by: NCI U01CA224012 and R01s CA186241, CA196228 and DoD PA210068. SB was supported by a postdoctoral fellowship from the German Research Foundation (BE 7224/1-1) and a research grant provided by the Sky Foundation. NGS is supported by NCI R00 CA263154 and the Henry Ford Health + Michigan State University Collaboration grant.

## References

1. Society, A.C., Cancer Facts & Figures 2024. American Cancer Society, 2024.

2. Chu, L.C., M.G. Goggins, and E.K. Fishman, Diagnosis and Detection of Pancreatic Cancer. The Cancer Journal, 2017. 23(6): p. 333–342.

3. Pihlak, R., J. Weaver, J. Valle, and M. McNamara, Advances in Molecular Profiling and Categorisation of Pancreatic Adenocarcinoma and the Implications for Therapy. Cancers, 2018. 10(1): p. 17.

4. Sahin, I.H., C.A. Iacobuzio-Donahue, and E.M. O’Reilly, Molecular signature of pancreatic adenocarcinoma: an insight from genotype to phenotype and challenges for targeted therapy. Expert Opinion on Therapeutic Targets, 2016. 20(3): p. 341–359.

5. Pancreatic Cancer: Molecular Characterization, Clonal Evolution and Cancer Stem Cells. Biomedicines, 2017. 5(4): p. 65.

6. Benitz, S., et al., ROR2 regulates cellular plasticity in pancreatic neoplasia and adenocarcinoma. 2023, Cold Spring Harbor Laboratory.

7. Schlesinger, Y., et al., Single-cell transcriptomes of pancreatic preinvasive lesions and cancer reveal acinar metaplastic cells’ heterogeneity. Nature Communications, 2020. 11(1).

8. Delgiorno, K.E., et al., Tuft Cells Inhibit Pancreatic Tumorigenesis in Mice by Producing Prostaglandin D2. Gastroenterology, 2020. 159(5): p. 1866–1881.e8.

9. Delgiorno, K.E., et al., Identification and Manipulation of Biliary Metaplasia in Pancreatic Tumors. Gastroenterology, 2014. 146(1): p. 233–244.e5.

10. Hoffman, M.T., et al., The Gustatory Sensory G-Protein GNAT3 Suppresses Pancreatic Cancer Progression in Mice. Cellular and Molecular Gastroenterology and Hepatology, 2021. 11(2): p. 349–369.

11. Bailey, J.M., et al., DCLK1 Marks a Morphologically Distinct Subpopulation of Cells With Stem Cell Properties in Preinvasive Pancreatic Cancer. Gastroenterology, 2014. 146(1): p. 245–256.

12. Nevo, S., N. Kadouri, and J. Abramson, Tuft cells: From the mucosa to the thymus. Immunology Letters, 2019. 210: p. 1–9.

13. Hoover, B., V. Baena, M.M. Kaelberer, F. Getaneh, S. Chinchilla, and D.V. Bohórquez, The intestinal tuft cell nanostructure in 3D. Scientific Reports, 2017. 7(1).

14. Howitt, M.R., et al., Tuft cells, taste-chemosensory cells, orchestrate parasite type 2 immunity in the gut. Science, 2016. 351(6279): p. 1329–1333.

15. Yamashita, J., M. Ohmoto, T. Yamaguchi, I. Matsumoto, and J. Hirota, Skn-1a/Pou2f3 functions as a master regulator to generate Trpm5-expressing chemosensory cells in mice. PLOS ONE, 2017. 12(12): p. e0189340.

16. Yamaguchi, T., et al., Skn-1a/Pou2f3 is required for the generation of Trpm5-expressing microvillous cells in the mouse main olfactory epithelium. BMC Neuroscience, 2014. 15(1): p. 13.

17. Ma, Z., et al., Single-Cell Transcriptomics Reveals a Conserved Metaplasia Program in Pancreatic Injury. Gastroenterology, 2022. 162(2): p. 604–620 e20.

18. Garcia, P.E., et al., Differential Contribution of Pancreatic Fibroblast Subsets to the Pancreatic Cancer Stroma. Cell Mol Gastroenterol Hepatol, 2020. 10(3): p. 581–599.

19. McGinty, J.W., H.-A. Ting, T.E. Billipp, M.S. Nadjsombati, D.M. Khan, N.A. Barrett, H.-E. Liang, I. Matsumoto, and J. Von Moltke, Tuft-Cell-Derived Leukotrienes Drive Rapid Anti-helminth Immunity in the Small Intestine but Are Dispensable for Anti-protist Immunity. Immunity, 2020. 52(3): p. 528–541.e7.

20. Matsumoto, I., M. Ohmoto, M. Narukawa, Y. Yoshihara, and K. Abe, Skn-1a (Pou2f3) specifies taste receptor cell lineage. Nature Neuroscience, 2011. 14(6): p. 685–687.

21. Hwang, W.L., et al., Single-nucleus and spatial transcriptome profiling of pancreatic cancer identifies multicellular dynamics associated with neoadjuvant treatment. Nature Genetics, 2022. 54(8): p. 1178–1191.

22. Caplan, L.R., V. Vavinskaya, D.G. Gelikman, N. Jyotsana, V.Q. Trinh, K.P. Olive, M.C.B. Tan, and K.E. Delgiorno, Enteroendocrine Cell Formation Is an Early Event in Pancreatic Tumorigenesis. Frontiers in Physiology, 2022. 13.

23. Farrell, A.S., et al., MYC regulates ductal-neuroendocrine lineage plasticity in pancreatic ductal adenocarcinoma associated with poor outcome and chemoresistance. Nature Communications, 2017. 8(1).

24. Wu, C.-Y.C., et al., PI3K Regulation of RAC1 Is Required for KRAS-Induced Pancreatic Tumorigenesis in Mice. Gastroenterology, 2014. 147(6): p. 1405–1416.e7.

25. Wen, H.-J., S. Gao, Y. Wang, M. Ray, M.A. Magnuson, C.V.E. Wright, M.P. Di Magliano, T.L. Frankel, and H.C. Crawford, Myeloid Cell-Derived HB-EGF Drives Tissue Recovery After Pancreatitis. Cellular and Molecular Gastroenterology and Hepatology, 2019. 8(2): p. 173–192.

26. Baine, M.K., et al., SCLC Subtypes Defined by ASCL1, NEUROD1, POU2F3, and YAP1: A Comprehensive Immunohistochemical and Histopathologic Characterization. Journal of Thoracic Oncology, 2020. 15(12): p. 1823–1835.

27. Huang, Y.-H., et al., POU2F3 is a master regulator of a tuft cell-like variant of small cell lung cancer. Genes & Development, 2018. 32(13-14): p. 915–928.

28. Madisen, L., et al., A robust and high-throughput Cre reporting and characterization system for the whole mouse brain. Nat Neurosci, 2010. 13(1): p. 133–40.

29. Burclaff, J., S.G. Willet, J.B. Sáenz, and J.C. Mills, Proliferation and Differentiation of Gastric Mucous Neck and Chief Cells During Homeostasis and Injury-induced Metaplasia. Gastroenterology, 2020. 158(3): p. 598–609.e5.

30. Westphalen, C.B., et al., Dclk1 Defines Quiescent Pancreatic Progenitors that Promote Injury-Induced Regeneration and Tumorigenesis. Cell Stem Cell, 2016. 18(4): p. 441–455.

31. Worthington, J.J., F. Reimann, and F.M. Gribble, Enteroendocrine cells-sensory sentinels of the intestinal environment and orchestrators of mucosal immunity. Mucosal Immunology, 2018. 11(1): p. 3–20.

32. Riedel, M.J., A. Asadi, R. Wang, Z. Ao, G.L. Warnock, and T.J. Kieffer, Immunohistochemical characterisation of cells co-producing insulin and glucagon in the developing human pancreas. Diabetologia, 2012. 55(2): p. 372–381.

33. Hu, L., et al., Genome-Wide Association Study of Prognosis in Advanced Non–Small Cell Lung Cancer Patients Receiving Platinum-Based Chemotherapy. Clinical Cancer Research, 2012. 18(19): p. 5507–5514.

34. Lim, U., et al., Susceptibility variants for obesity and colorectal cancer risk: The multiethnic cohort and PAGE studies. International Journal of Cancer, 2012. 131(6): p. E1038–E1043.

35. Ireland, A.S., et al., MYC Drives Temporal Evolution of Small Cell Lung Cancer Subtypes by Reprogramming Neuroendocrine Fate. Cancer Cell, 2020. 38(1): p. 60–78.e12.

36. Lee, J.K., et al., N-Myc Drives Neuroendocrine Prostate Cancer Initiated from Human Prostate Epithelial Cells. Cancer Cell, 2016. 29(4): p. 536–547.

37. Mollaoglu, G., et al., MYC Drives Progression of Small Cell Lung Cancer to a Variant Neuroendocrine Subtype with Vulnerability to Aurora Kinase Inhibition. Cancer Cell, 2017. 31(2): p. 270–285.

38. Yin, Y., et al., N-Myc promotes therapeutic resistance development of neuroendocrine prostate cancer by differentially regulating miR-421/ATM pathway. Molecular Cancer, 2019. 18(1).

39. Barker, N., Adult intestinal stem cells: critical drivers of epithelial homeostasis and regeneration. Nature Reviews Molecular Cell Biology, 2014. 15(1): p. 19–33.

40. Sinha, S., et al., PanIN Neuroendocrine Cells Promote Tumorigenesis via Neuronal Cross-talk. Cancer Research, 2017. 77(8): p. 1868–1879.

41. Koga, H., Y. Ikezono, and T. Torimura, Pancreatic DCLK1 marks quiescent but oncogenic progenitors: a possible link to neuroendocrine tumors. Stem Cell Investigation, 2016. 3: p. 37–37.

42. Nakanishi, Y., et al., Dclk1 distinguishes between tumor and normal stem cells in the intestine. Nature Genetics, 2013. 45(1): p. 98–103.

43. Saqui-Salces, M., T.M. Keeley, A.S. Grosse, X.T. Qiao, M. El-Zaatari, D.L. Gumucio, L.C. Samuelson, and J.L. Merchant, Gastric tuft cells express DCLK1 and are expanded in hyperplasia. Histochemistry and Cell Biology, 2011. 136(2): p. 191–204.

44. Nawabi, H., et al., Doublecortin-Like Kinases Promote Neuronal Survival and Induce Growth Cone Reformation via Distinct Mechanisms. Neuron, 2015. 88(4): p. 704–719.

45. May, R., et al., Identification of a novel putative pancreatic stem/progenitor cell marker DCAMKL-1 in normal mouse pancreas. Am J Physiol Gastrointest Liver Physiol, 2010. 299(2): p. G303–10.

46. Xu, L., M. Zhang, L. Shi, X. Yang, L. Chen, N. Cao, A. Lei, and Y. Cao, Neural stemness contributes to cell tumorigenicity. Cell &amp; Bioscience, 2021. 11(1).

47. Zhang, Z., et al., Similarity in gene-regulatory networks suggests that cancer cells share characteristics of embryonic neural cells. Journal of Biological Chemistry, 2017. 292(31): p. 12842–12859.

48. Beltran, H., et al., The role of lineage plasticity in prostate cancer therapy resistance. Clinical Cancer Research, 2019: p. clincanres.1423.

49. Dai, M.-S., Y. Jin, J.R. Gallegos, and H. Lu, Balance of Yin and Yang: Ubiquitylation-Mediated Regulation of p53 and c-Myc. Neoplasia, 2006. 8(8): p. 630–644.

50. Feng, Y.C., et al., c-Myc inactivation of p53 through the pan-cancer lncRNA MILIP drives cancer pathogenesis. Nature Communications, 2020. 11(1).

51. Ho, J.S.L., W. Ma, D.Y.L. Mao, and S. Benchimol, p53-Dependent Transcriptional Repression of c-myc Is Required for G1 Cell Cycle Arrest. Molecular and Cellular Biology, 2005. 25(17): p. 7423–7431.

52. Wu, J., et al., Generation of a pancreatic cancer model using a Pdx1-Flp recombinase knock- in allele. PLOS ONE, 2017. 12(9): p. e0184984.

53. Zhang, H., et al., Glucose-Mediated Repression of Menin Promotes Pancreatic β-Cell Proliferation. Endocrinology, 2012. 153(2): p. 602–611.

