## Supplementary for "Tuft cells transdifferentiate to neural-like progenitor cells in the progression of pancreatic cancer"

***Supplementary Table 1:*** Antibodies Used for Immunohistochemistry and Immunofluorescence.

| Antibody | Company | Catalog# | Dilution | Species |
|---|---|---|---|---|
| tdTomato (tdTom) | LifeSpan Bio | LS-C340696 | 1:250 (IF); 1:1500 (IHC) | Anti-Goat |
| Pou2f3 | Santa Cruz Biotechnology | sc-330 | 1:100 | Anti-Rabbit |
| Dclk1 | Abcam | ab31704 | 1:1000 (IF&IHC) | Anti-Rabbit |
| Vav1 | Cell Signaling | 2502s | 1:200 | Anti-Rabbit |
| Cox1 | Invitrogen | MA5-32259 | 1:1000 | Anti-Rabbit |
| Synaptophysin (Syp) | Sigma | 336R-95 | 1:200 (IF/IHC) | Anti-Rabbit |
| Gkn1 | Invitrogen | PA5-47913 | 1:100 | Anti-Sheep |
| Lectin GS-II | Invitrogen | L21415 | 1:100 | 488 Conjugated |
| Glucagon (Gcg) | Cell Signaling | 2760s | 1:100 | Anti-Rabbit |
| Insulin (Ins) | Novus Biological | MAB1417 | 1:150 | Anti-Rat |
| Somatostatin (Sst) | Phoenix Pharmaceuticals | h-060-03 | 1:100 | Anti-Guinea Pig |
| Ghrelin (Ghrl) | Cell Signaling | 31865s | 1:100 | Anti-Rabbit |
| Pancreatic Polypeptide (Pp/Ppy) | Abcam | ab272732 | 1:200 | Anti-Rabbit |
| Serotonin (5-HT) | Immunostar | 20080 | 1:250 | Anti-Rabbit |
| Nrxn3 | Alomone Labs | ANR-033 | 1:100 | Anti-Rabbit |
| Acetylated alpha Tubulin | Sigma | T7451-100UL | 1:1000 | Anti-Mouse |
| Wide-spectrum cytokeratin (PanCK) | Abcam | ab9377 | 1:500 | Anti-Rabbit |

**Supplementary Figure 1: Specific expression of tdTomato in Tuft cells**

IF is used to determine specificity of tdTomato reporter with tuft cells using different markers of tuft cells in *KF-P2f3T* **(A)** and *KPF-P2f3T* **(B)**. Cox1, Dclk1, Vav1 and Trpm5 in Green are co-stained with tdTom (Red) and Acetylated-α-Tubulin (Magenta). Scale bars = 10 μm unless otherwise noted.

**Supplementary Figure 2: *KPF-P2f3T* mTCS do not express Gastric Pit-like or Gastric Chief-like Markers.**

Characterization of mTCs and mNECs in neoplasia and Carcinoma models. **A)** Gkn1 (Gastric Pit-like Marker), Lectin II (Gastric Neck-like Marker), and Syp (Neuroendocrine Marker) in Green co-stained with tdTom (Red) and Hoechst (White) for both *KF-P2f3T* and *KPF-P2f3T* models. **B)** tdTom (Red) and Insulin (Green) are co-stained in *KF-P2f3T* and *KPF-P2f3T* models to investigate tdTom expression in islets. **C)** KPCY model was stained to demonstrate that mTCs stained by Cox1 (Red) and mNECs stained by Syp (Magenta) are derived from acinar cells via Ptf1a-Cre driven YFP (Green) in mice as in KCY models in previous studies. Scale bars = 10 μm unless otherwise noted.

**Supplementary Figure 3: mTCs do not require proliferation to transdifferentiate into mNECs.**

**A)** KF; Pou2f3$^{CreERT/CreERT}$ and **B)** KPF; Pou2f3$^{CreERT/CreERT}$ genetic strategy to deplete tuft cell populations. **C)** EdU (Magenta) was used to mark proliferating cells post-tamoxifen treatment when mTCs are known to transdifferentiate into mNECs. Cox1 (Green), tdTom (Red) Syp (White) and Hoechst (Blue) were co-stained to determine proliferation of cells (Magenta arrows). The yellow arrow indicates transitionary cell marked with Cox1, tdTom, and Syp. **D)** Co-stained merged images of islets and lesions from *KF-P2f3T* and *KPF-P2f3T* for Serotonin (5-HT) in Green with Syp (Red), PanCK (Teal), and Hoechst (White) to confirm lack of β cell identity from mNECs in PDA (*Left*).

Individual channels of merged images in Figure 3B (*Right*). Scale bars = 10 µm unless otherwise noted.

**Supplementary Figure 4: Targeted cMyc expression to modulate Tuft-to-NRP-Transition**

KF-MOE **(A)** and KPF-P2M$^{Fl/+}$ **(B)** genetic strategy to modulate cMyc expression specifically in tuft cells as PDA progresses. **C)** *KF-MOE* at 7 weeks post-tamoxifen treatment are stained with tdTom (Red), Syp (Green), Acetylated-α-tubulin (Magenta), and Hoechst (White) to identify mTC-derived mNECs in both time points. **D)** Co-staining of Syp (Green), tdTom (Red), PanCK (Magenta), and Hoechst (White) in KPF-P2M$^{Fl/+}$ at 7 weeks post-tamoxifen treatment to identify Syp+ cells and tdTom+ cells are separate populations when you knock down *cMyc* expression. Scale bars = 10 µm unless otherwise noted.

### Supplementary Figure 1

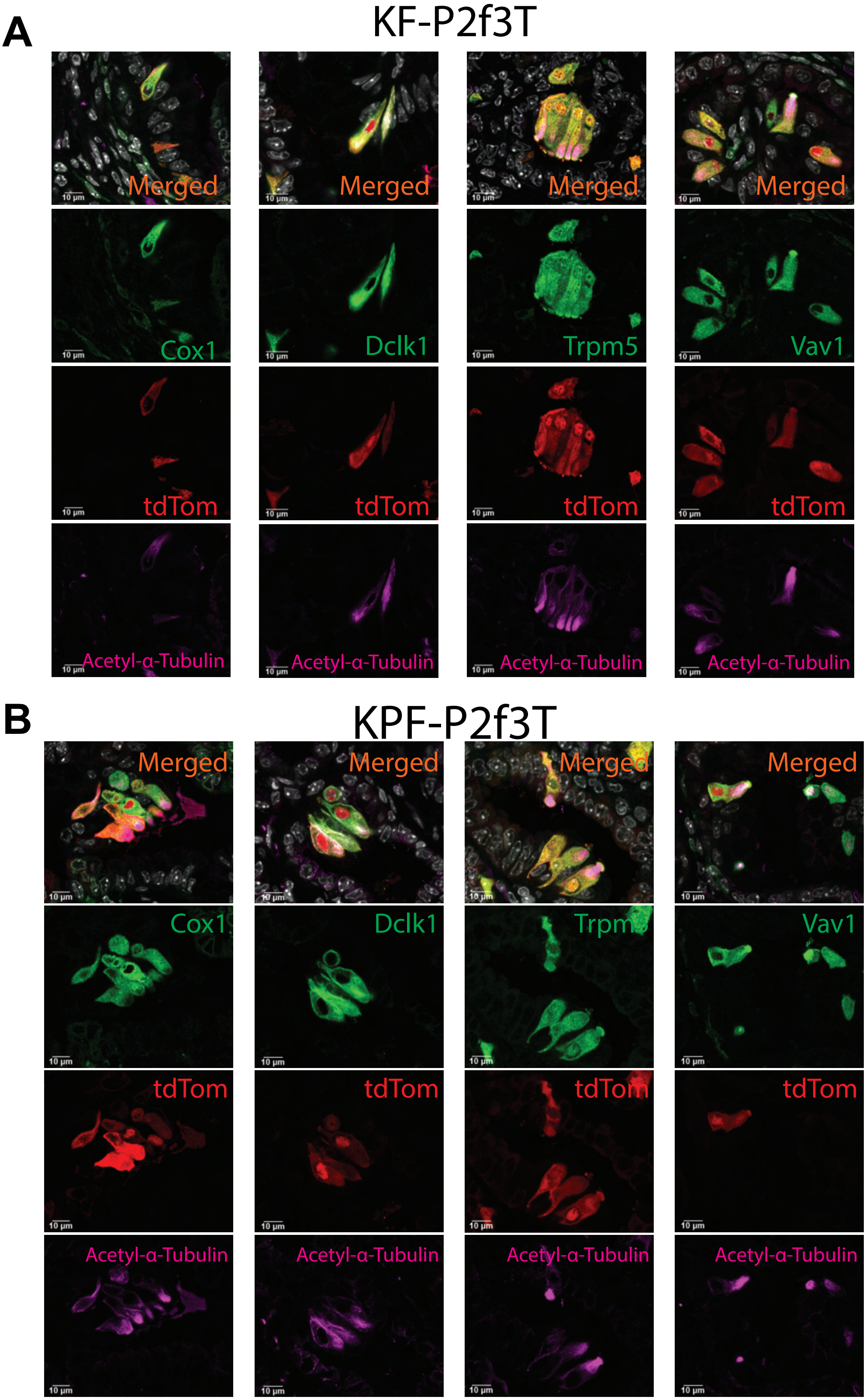

### Supplementary Figure 2

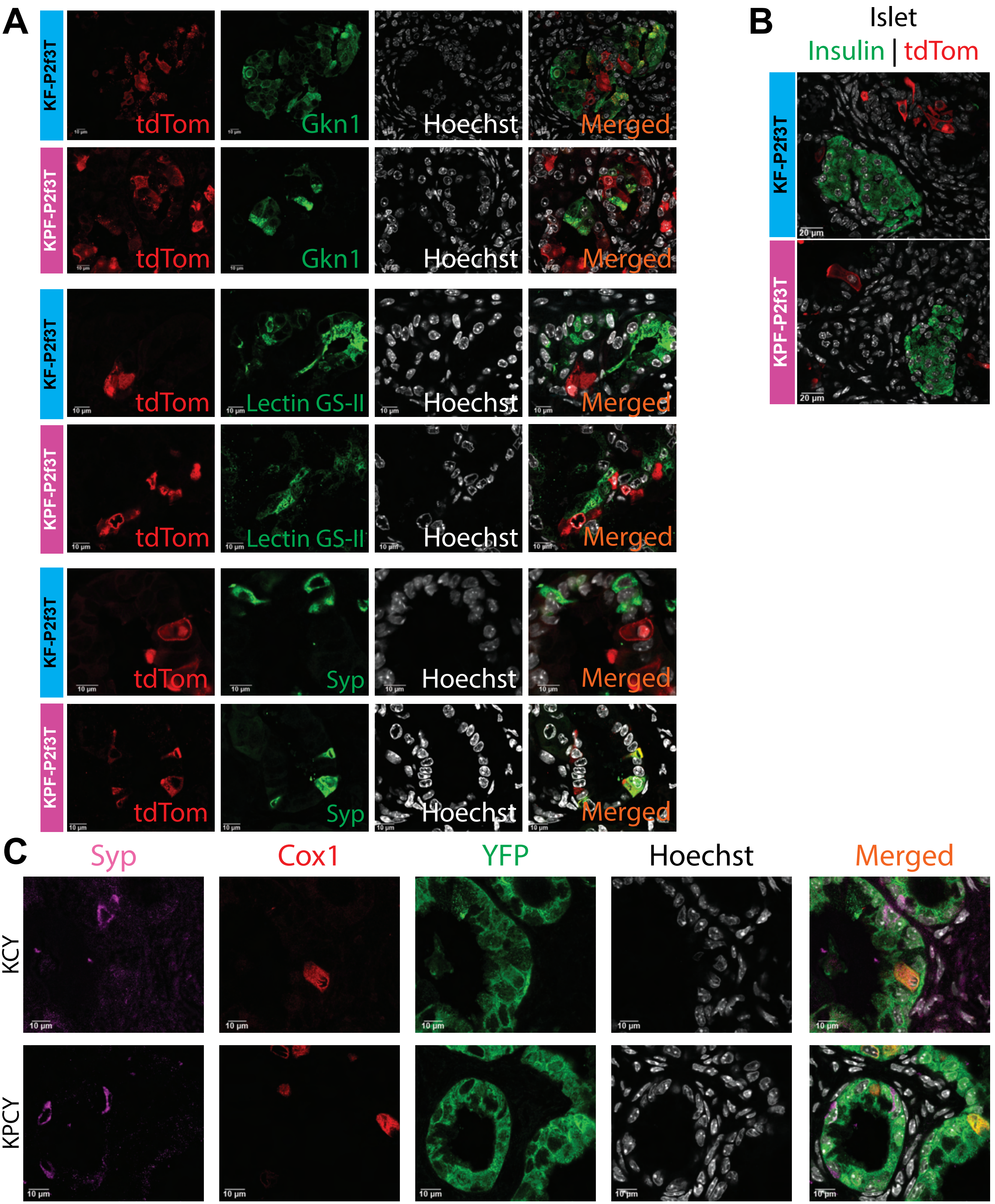

### Supplementary Figure 3

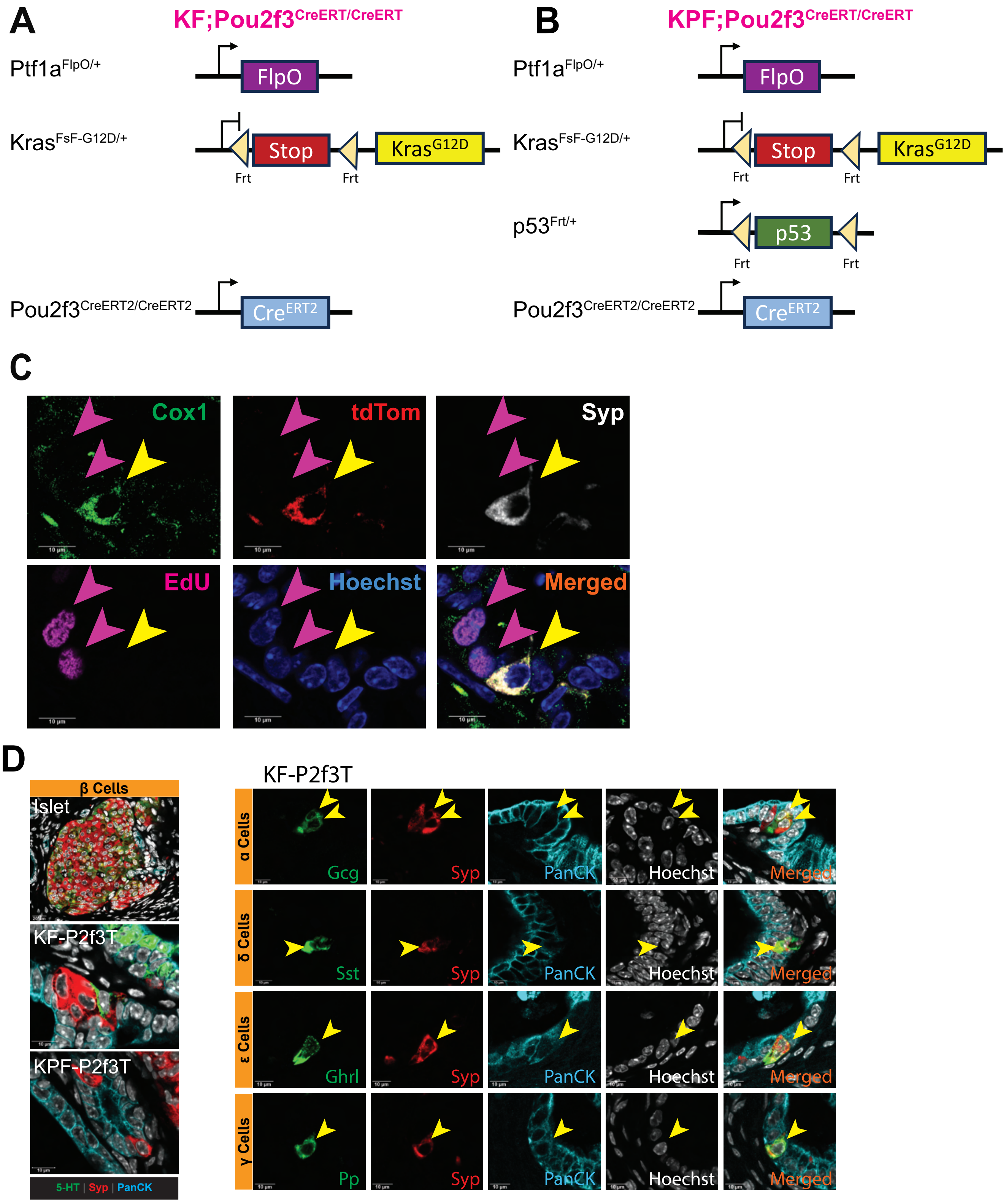

### Supplementary Figure 4

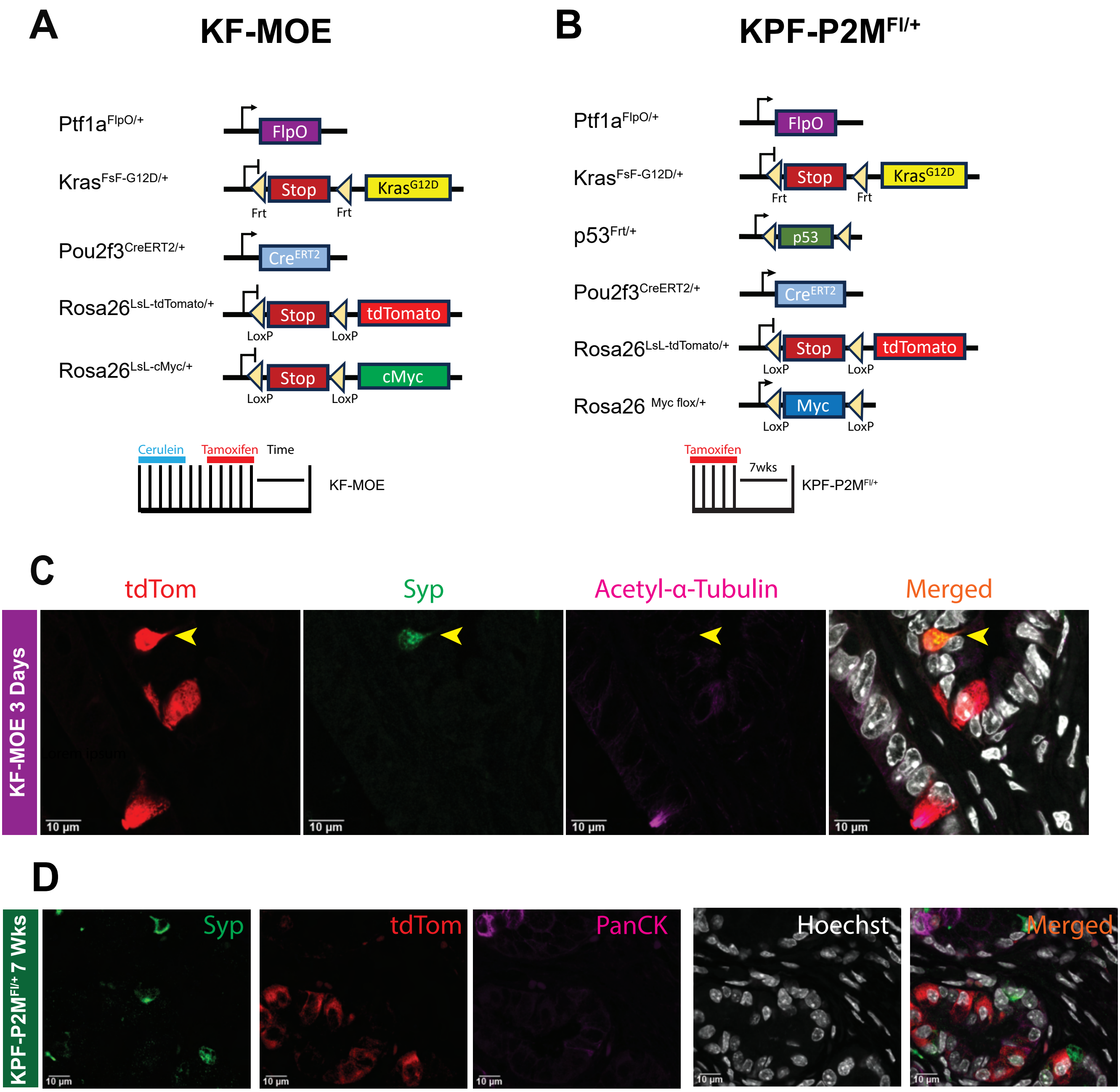
